# synpact: accurate, memory-light PacBio HiFi read mapping via a hierarchy of locally-consistent syncmer blocks

**DOI:** 10.64898/2026.06.28.735066

**Authors:** Mahmud Sami Aydın, Kristoffer Sahlin

## Abstract

**Motivation:** Mapping PacBio HiFi reads is a routine task and serves as a central step in many bioinformatics analyses. However, the most accurate long-read mappers have a high memory consumption and are slow. Some light-weight mappers have been proposed for faster runtime, but their accuracy is not comparable to state-of-the-art mappers. With the increasing number of available reference sequences, memory-efficient and fast methods for read mapping without the large accuracy drop are desired. A general trade-off with seed-chain-extend mappers is selecting a single, fixed seed size, which forces a compromise between sensitivity and specificity.

**Results:** We present `synpact`, a long-read mapper that uses several seed sizes (a *hierarchy*) constructed with Locally Consistent Parsing (LCP) over syncmers. A read is mapped by querying for matches at different levels, followed by sliding window voting. By storing only the coarse upper levels rather than the full hierarchy, the index holds several times fewer entries, while still handling errors by falling back from coarser to finer stored levels at query time. We benchmark `synpact` against popular long-read mappers on four genomes and different read lengths. For simulated PacBio HiFi data, `synpact` matches or approaches `minimap2` accuracy with higher precision in most cases, while using roughly 5–13× less peak memory (e.g., ∼0.8 GB vs. ∼10.7 GB on human) and mapping faster on large or repetitive genomes (e.g., about 10 to 13 times faster than `minimap2` on rye). On real HiFi reads `synpact` has high concordance with `minimap2` across the four genomes, as opposed to the other lightweight long-read mappers. Availability and Implementation: `synpact` is written in Rust and is available at https://github.com/mahmudsami/synpact

## 1. Introduction

PacBio HiFi reads now have a per-base accuracy of about 99.8 % [28], and have become standard input for genome assembly, structural variant discovery, and pangenome construction. The first step in many long-read pipelines is read mapping, i.e., finding the origin of each read in a reference genome. At HiFi error rates, the challenge is no longer handling sequencing errors, but resolving reference ambiguity from segmental duplications, paralogous gene families, and tandem repeats [4], while keeping runtime and memory usage manageable for large sequencing datasets.

Many long-read mapping algorithms have been developed [24]. The most widely used long-read mapper, `minimap2` [17], selects minimizer seeds, chains them, and extends chains with base-level alignment. It achieves high mapping accuracy, but requires several gigabytes of memory for a human genome index, and both runtime and memory scale poorly on large repetitive genomes such as many plant genomes. There are also other long-read mappers with different design goals. For example, `NGMLR` [25] was developed specifically to improve mapping sensitivity around structural variants, and `Winnowmap2` [16] aims to achieve accurate mapping in repetitive regions by decreasing the weight of high-frequency minimizers, but is slower than minimap2 [13]. Finally, there is a category of long-read mappers designed for fast mapping PacBio HiFi reads that use longer fuzzy seeds and often trade accuracy for lower resource usage. BLEND [13] extends minimizers to allow fuzzy long seed matches through strobemers [22] and is fast and memory-efficient, while `mapquik` [12] constructs long *k*-min-mer seeds for fast mapping. However, as we show, their accuracy decreases on large, repeat-rich genomes, or when the error or mutation rate increases.

Instead of using only short or long seeds, we argue that HiFi mapping benefits from a *hierarchy* of seeds at different levels. The key observation is that, at HiFi error rates, one can often afford to map reads with longer seeds as they are rarely corrupted by errors, but in case of errors or mutations, falling back to shorter seeds is useful to increase sensitivity. Such a hierarchy has been explored in the read mapper strobealign [23] through multi-context-seeds [27]. However, similar to minimap2, strobealign is accurate, but requires several gigabytes of memory for a human genome index, and is not yet designed for long-read mapping [27].

Locally consistent parsing (LCP) is an approach to segmenting sequences into *blocks* (i.e., segments) such that most block boundaries are preserved under small edits [21]. It has been used in sequence compression [15], whole genome alignment [1], genome distance estimation [2], and its theoretical properties have been thoroughly explored [7, 6]. Unlike minimizers, which select a single *k*-mer per window, LCP methods partition the entire sequence into partially overlapping blocks based on local sequence content. A key property is that a single substitution or indel affects only the blocks in its immediate vicinity, leaving the rest of the parse unchanged. This stability under errors, and its different hierarchies, makes LCP an attractive foundation for read mapping, where the goal is to find matching blocks between a read and a reference despite a small number of differences.

In this paper, we propose a new resource-frugal read mapper, `synpact`, that combines syncmer sketching (for initial sequence reduction) with the hierarchical parsing of LCP to create seeds for read mapping at different resolution levels (hierarchy). By storing only some of the levels, `synpact`’s index is roughly an order of magnitude smaller than that of, e.g, minimap2. `synpact` first searches for seed matches at the highest levels (longest seeds), but visits lower levels if needed, for higher granularity in regions with errors or mutations. After finding seeds at various resolutions, `synpact` then uses a weighted seed voting to find candidate mapping loci.

`synpact` was developed specifically to improve accuracy and resource usage over state-of-the-art in the lightweight long-read mapper class among tools such as BLEND and `mapquik`, rather than to further push the accuracy ceiling of full-alignment tools like `minimap2`, `Winnowmap2` [16], or `NGMLR`. We benchmark `synpact` against `minimap2`, BLEND, and `mapquik` on simulated and real HiFi data across four genomes, showing it matches or approaches the accuracy of minimap2 at a fraction of the memory and at competitive or better speed. Among the lightweight read-mappers (BLEND, mapquik, `synpact`), `synpact` is the only one of these three that remains accurate on large, repeat-rich genomes, and it uses the least memory of any mapper in the comparison. We further demonstrate the robustness of `synpact` by mapping simulated reads to the 36Gb lungfish genome at higher accuracy, lower runtime, and lower memory usage than minimap2 (with a sparse index), showing the benefit of `synpact`’s hierarchical approach beyond just creating a sparser index.

## 2. Methods

`synpact` turns each sequence (reference or read) into a multi-level hierarchy of seeds, referred to as *blocks* below, in the context of LCP. The hierarchical LCP block encoding of syncmers (described in sections 2.1-2.3) is applied to the reference to form the index (section 2.4). The blocks are also constructed from the reads, which are queried against the index and serve as anchors. The anchors are then grouped by diagonal voting (section 2.5). `synpact` then outputs the mapping locations in PAF format.

### 2.1. Open-syncmer *k*-mer sampling

Let Σ = {A, C, G, T} and let *S* ∈ Σ^∗^ be a reference contig or a read. We sketch *S* with *open syncmers* [11] as follows. Fix a *k*-mer length *k* and an *s*-mer length *s < k* (defaults *k* = 19, *s* = 10) and a hash function *h* : Σ^*s*^ →Z on *s*-mers. A *k*-mer *K* = *S*[*i* .. *i*+*k*−1] contains the *k* − *s* + 1 *s*-mers *K*_*j*_ = *K*[*j* .. *j*+*s*−1] for 0 ≤ *j* ≤ *k* − *s*. Let

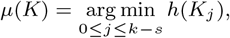

i.e., the offset of its minimum-hash *s*-mer, *K* is selected as an *open syncmer* iff *µ*(*K*) = *t* for a fixed offset *t*; we use the central offset *t* = ⌊(*k* − *s*)*/*2⌋, which maximises the regularity of inter-seed spacing and the conservation of selected *k*-mers under substitutions, as described in [11]. Under a uniform hash every offset is equally likely, so the expected density of selected *k*-mers is 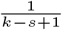.

Two properties make this sketch suitable for mapping. First, the syncmer selection is *local* (i.e., context independent), which means that *µ*(*K*) depends only on the relative order of the *s*-mer hashes inside *K*. This makes syncmers more *error-tolerant* than minimizers (given the same size of the seed). Secondly, syncmers lower-bounds the minimum distance between seeds, in our case to *t* + 1, which can benefit mapping. However, it should be noted that closed syncmers and minimizers, unlike open syncmers, provide upper bound distances between consecutive seeds. Nevertheless, we have found that open syncmers yield slightly favorable results over the other two seeding approaches. Both strands of the read are sketched by processing *S* and its reverse complement as two independent passes.

### 2.2. Locally consistent parsing into blocks

Originally, the Locally Consistent Parsing (LCP) scheme, described in [21], partitions a string over an ordered alphabet into short, non-overlapping substrings, called *blocks* [7] (sometimes also called *cores* [1]), so that (i) equal substrings are partitioned identically up to their margins (*partition consistency*) and (ii) every symbol belongs to some block (*contiguity*) [21].

Rather than processing the entire sequence with LCP directly, we parse the syncmers into blocks (i.e., groups of syncmers). Treating the syncmer hash values (64 bits) as the alphabet, and the sequence of syncmer hash values *v*_1_*v*_2_ · · · as the input string over the value order *<*. A *position* is a block boundary according to the four block-identification rules, described in LCPan [1], that we state here for completeness. For a substring of a current string, we write the relevant substring as … *x y z* … for rule 1 and 2, and as … *x y*^*i*^ *z* … for rule 3:

1. **Local minimum (LMIN)**. A length-3 substring *xyz* is a block when its middle hash value is a strict local minimum, *x > y < z*.
2. **Local maximum (LMAX)**. A substring *xyz* is a block when its middle hash value is a strict local maximum, *x < y > z*, and neither neighboring hash value is itself a local minimum (so an LMAX block is not declared adjacent to an LMIN block).
3. **Repetitive run (RINT)**. A maximal run *x y* ^*i*^ *z* with *i >* 1, *x* ≠ *y* and *y* ≠ *z*, i.e., a stretch of one repeated value bounded by two distinct hash values, forms one block.
4. **Monotone run (SSEQ)**. A maximal strictly increasing or strictly decreasing run that links two neighboring blocks (its endpoints belonging to LMIN/LMAX/RINT blocks) forms a block.

An illustration of the four rules are shown in Figure 1a. This rule set upholds the partition consistency criterion, since local minimum and maximum, repetitions, and monotonic sequences are not affected by distant sequence mutations. Also, a hash value must belong to at least one of the four categories LMIN, LMAX, RINT, SSEQ. Thus, it is contiguous as well. In [1], the authors show that blocks cannot start at three consecutive positions in any level, therefore the number of blocks are bounded. However, the four rules by themselves do not guarantee a bounded-size block (e.g., a monotone infinite sequence), and may thus produce an unbounded block size which is detrimental for read mapping. Thus, we modify the fourth rule to bound its block size using Deterministic Coin Tossing (DCT) [10] in a different fashion to LCPan. We discuss this LCP modification in detail in Suppl. Section 1.

**Figure 1.**
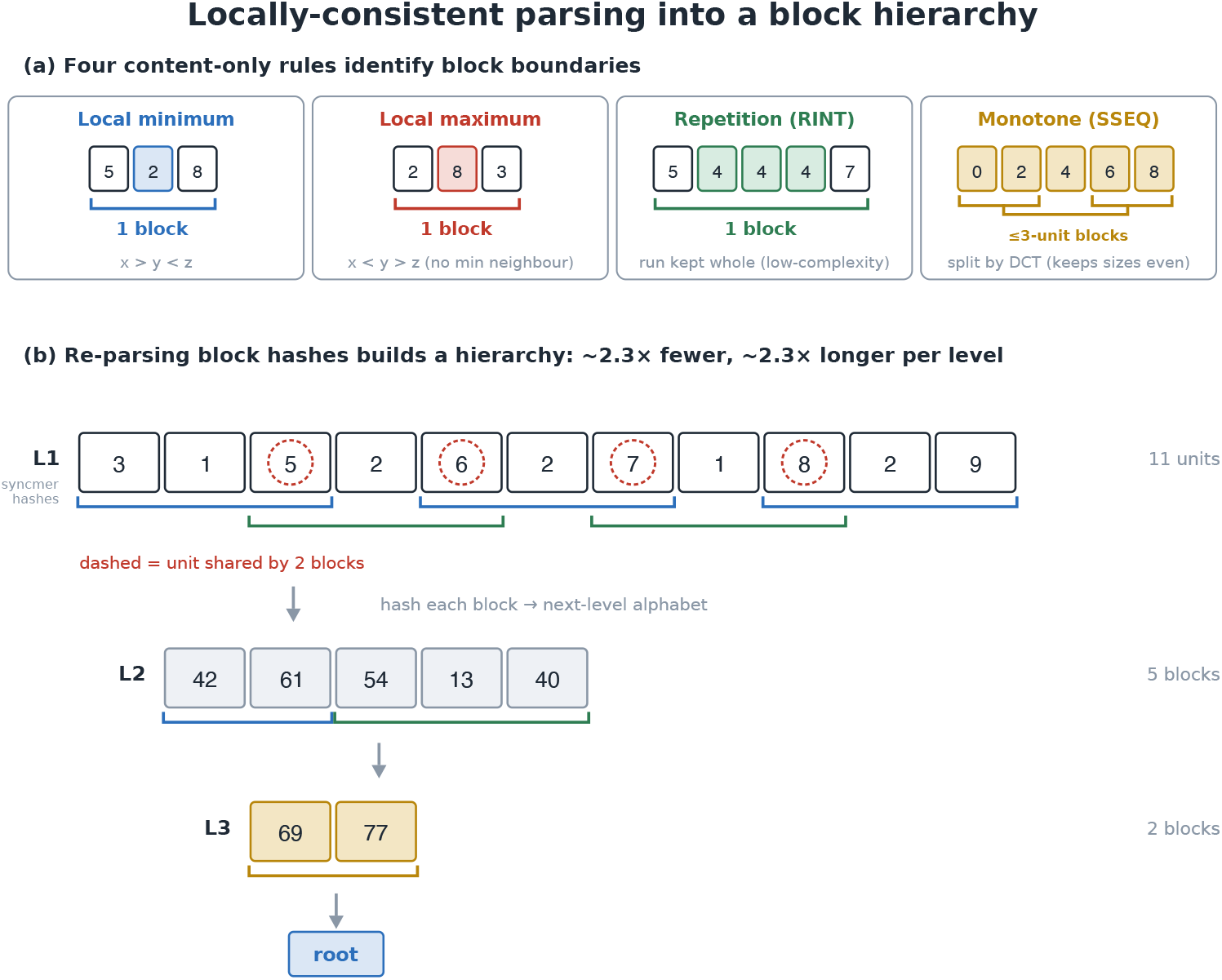
Locally-consistent parsing and the block hierarchy. (a) The four content-only rules that mark block boundaries: local minimum, local maximum, repetition kept whole (i.e., a low-complexity region), and monotone runs split by DCT into ≤ 3-unit blocks. (b) Re-parsing block hashes builds a hierarchy with ∼2.3× fewer and ∼2.3× longer blocks per level. A lower-level block shared between adjacent blocks (red circles).

### 2.3. Recursive block hierarchy

Applying the parse over syncmers described in the previous section forms the first *level* of seeds (L1). Each block is represented by a 64-bit hash value formed by a shifted XOR operation of the syncmer hash values. The same LCP rules are applied recursively to the sequence of L1 block hash values to form L2 blocks, and to L2 blocks to form L3 blocks, etc, up to six levels. Figure 1b illustrates the concept. Unlike LCPan [1], we do not apply a deterministic coin-tossing between levels. The 64-bit block hashes already form a large, content-determined alphabet, so parsing them directly keeps the parse locally consistent. That is, identical block-hash subsequences on the read and the reference yield identical higher-level blocks. Each iteration shrinks the number of blocks and lengthens them by a near-constant factor (measured to *c* ≈ 2.3 [1]). This means the blocks are on average 37 bp at L1, about 188 bp in L3, and about 2.2 kb at L6 (measured empirically on the human CHM13 assembly). So a read may be anchored by its few high-level blocks, which are long enough to be unique on the genome, but a long block is also more likely to contain a sequencing error. If a high-level block is not found in the index, the read falls back to shorter blocks, illustrated in Figure 2.

**Figure 2.**
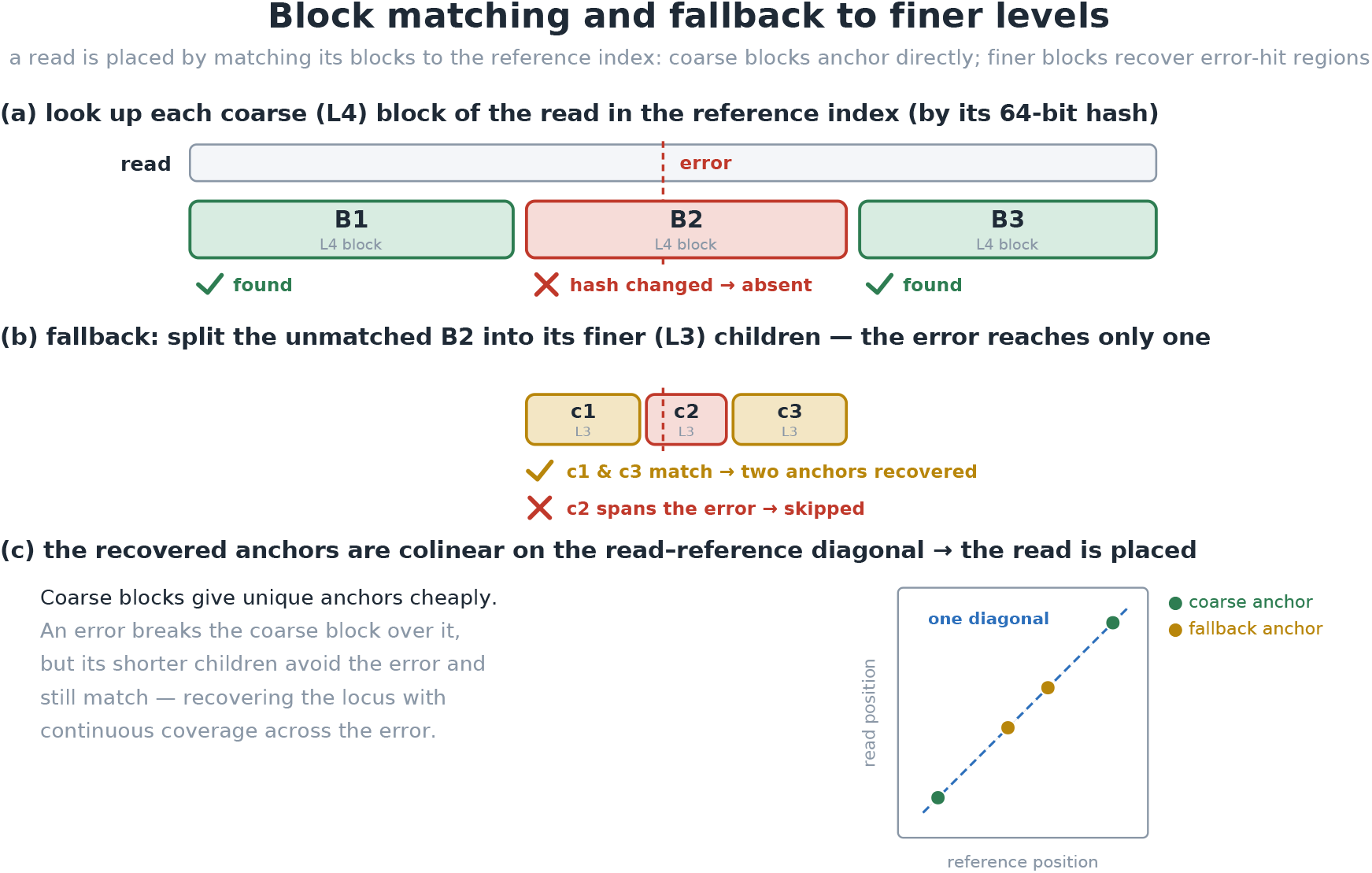
Block matching and fallback to finer levels. A toy example with two levesl L1 and L2. (a) Each coarse (L2) block of a read is looked up in the reference index by its hash value. B1 and B3 are found in the index, while the block spanning the error is not found. (b) That block falls back to its children blocks c1, c2, and c3 in L1. (c) The recovered anchors are colinear on the read–reference diagonal, so the read is placed with continuous coverage across the error.

### 2.4. Reference index

For every indexed level, the reference index stores three parallel arrays `(hash, chr, pos)` for the hash value (8 bytes), reference identifier (1 byte), and position (4 bytes) of each block (acting as a seed for mapping). The arrays are sorted by the hash value. We represent this as a struct-of-arrays layout (separate arrays) rather than an array-of-structs, which would require padding the remaining 3 bytes for an even 16 bytes. Also, the struct-of-arrays only requires binary search over the array of hashes (8 bytes), thus fitting more elements for the same size of cache.

### 2.5. Anchoring and diagonal voting

A read is encoded into blocks at different levels, in the same manner as the reference sequences (as described in previous sections), on both strands and matched against the index, starting with higher levels. Each block is queried by its hash value. If a block is absent, it is replaced by querying the children blocks at a lower level (Figure 2). Only levels at or above the index floor (min-level, default 3) are queried. Every successful query yields an *anchor* with the information (*chr, q*_pos_, *r*_pos_, *w*) at the matched reference position, where *chr* is reference chromosome, *q*_pos_ is read position and *r*_pos_ is reference position. The anchor weight *w* is defined as the number of syncmers within the anchor. If a syncmer is contained by multiple seeds, its weight is divided between the blocks. Since the blocks are only sharing syncmers in the ends, the possible weights of a syncmer in an anchor are 1, 1/2, and (in rare cases) 1/3.

The read’s locus is then chosen by *diagonal voting* (details in Suppl. section 2). Briefly, anchors are sorted by chromosome and diagonal *d* = *r*_pos_ − *q*_pos_. A window of size *W* sweeps over the diagonal vector and computes the total syncmer weight for each eligible window. The heaviest window is the read’s assigned locus. Disjoint loci, i.e., loci on different chromosomes or on a diagonal more than *W* away, give alternative candidate loci. For the primary locus of weight (*w*_*P*_) and second *w*_*S*_ loci, the mapping quality is then computed as `mapq` = min(60, ⌊(*w*_*P*_ −*w*_*S*_)·60*/w*_*P*_ ⌋), so a read with one dominant locus scores 60 and one with two equally-supported loci scores near 0. Sorting the anchors dominates the time complexity, making placement *O*(*n* log *n*) in the number of anchors.

### 2.6. Implementation details

We use *k* = 19 and *s* = 10 for the open syncmers as default, which corresponds to an expectation of one seed per 10 positions. We index only blocks from levels 3 to 6 (parameters; default min-level is 3, and max-level 6). This setting works well for a uniform error rate of about 1%, or potentially higher error rates if errors are clustered (as can happen in real sequencing data. We set the window size W to a default of 500 bp.

## 3. Results

`synpact` is designed to be a resource-light mapper and compete with other lightweight read mappers in that class (e.g., mapquik, BLEND). We also include the popular, accurate but resource-heavy, `minimap2` in the benchmark as a proxy for state-of-the-art long-read mapping accuracy. We aimed to assess whether `synpact` can be more robust and achieve higher accuracy than other lightweight mappers when mapping HiFi data, while requiring much less resources compared to `minimap2`.

### 3.1. Experimental setup

We benchmarked `synpact` (k=19, s=10, min-level=3) against `minimap2` (`-x map-hifi –secondary=no`), BLEND and `mapquik` on five genomes spanning a range of size and complexity (see Table 1). We also use the CHM13 chromosome Y (*human_y*; `NC_060948.1`), as an additional dataset since it is a small but highly repetitive ampliconic/satellite sequence for a targeted test of repeat disambiguation. We ran all experiments on a MacBook Pro with Apple M2 Max and 32 GB RAM with an Ubuntu:22.04 docker container.

**Table 1.** Genomes and HiFi read datasets used in the benchmarks. Each read set was subsampled to the first 100,000 reads. Reads come from the same reference as from which the assembly is built, except for human, where HG002 reads are mapped to the CHM13 reference (cross-individual, ∼0.1 % divergence).

| Genome | Reference identifier (citation) | Size | Platform | Accession |
| --- | --- | --- | --- | --- |
| *Arabidopsis* | Col-CEN v1.2 [19] | 131 Mb | Col-0 HiFi | ERR13987671 |
| Maize | Mo17 T2T [8] | 2.2 Gb | Mo17 HiFi | SRR15447414 |
| Human | T2T-CHM13v2.0 [20] | 3.1 Gb | HG002 Revio HiFi (HPRC) | SRR34290932 |
| Rye | Lo7_V3 [26] | 7.9 Gb | Lo7 Revio HiFi | ERR15194059 |
| Rye | Lo7_V3 [26] | 7.9 Gb | Lo7 Revio (DeepConsensus) | ERR15194060 |
| Lungfish | neoFor_v3 [18] | 36 Gb | Simulated | Simulated |

#### Simulated reads

For each genome, we generated 100,000 simulated HiFi reads at four error rates (0, 0.1, 0.5 and 1 %) and four mean read lengths (10, 15, 20, and 25 kb) with standard deviations of (500, 750, 1000, 1250), respectively. Reads were drawn from random reference loci with 50% substitutions, 25% insertions, and 25% deletions. A read is counted as *correct* if its reported start lies within 1,000 bp of its true start. We report *mapping rate* (fraction of mapped reads), *accuracy* (fraction of *all* reads placed correctly), *precision* (fraction of *mapped* reads that are correct), and wrong-chromosome calls.

#### Real reads

In the absence of ground truth, we assessed real read mapping performance using mapping agreement. A locus is accepted as truth only where at least two mappers agree on the chromosome within at most 1000 bp difference in starting position. While this is not as precise as having known read locations, it serves as a reasonable proxy and primarily suffices to clearly separate read mapper accuracy, as we will show. We mapped subsets consisting of the first 100,000 reads from PacBio HiFi read datasets (Table 1) against each respective reference.

We included two versions of the reads from the rye genome (regular HiFi and a DeepConsensus readset [3]). DeepConsensus re-polishes HiFi CCS reads to make them substantially more accurate [3]. It therefore provides nearly error-free data that allows us to compare performance on real reads with various error rates (unpolished vs polished) within the ≤ 1 %-error range `synpact` is designed for.

#### The lungfish genome stress test

To demonstrate `synpact`’s performance and improved scaling over the other methods beyond the four reference genomes, we also included a benchmark on the ∼36 Gb Australian lungfish genome (*Neoceratodus forsteri*), which we refer to as *lungfish*. The lungfish genome is roughly an order of magnitude larger than human and among the largest animal genomes assembled. At this size a default `minimap2` using the `map-hifi` setting, as well as the other mappers, are not possible to run on our machine due to RAM constaints. Instead we compare `synpact` against a *sparsified* `minimap2` setting (maximal *k* = 28 and a much coarser minimizer window *w* = 200). On this genome we compared `synpact` to `minimap2` across the same four error rates and four read lengths as for the other simulated data.

#### Time and memory

We measured time and peak memory for index construction and read mapping separately for all of our experiments. For each tool the index is built once per genome, recording its build time and peak memory usage. The reads are then mapped against that pre-built index. `mapquik` is the exception, as it has no separate indexing step. All tools are run with eight threads. However, for `synpact` we additionally record a single-threaded index build alongside the default eight-thread build, to report index-construction tradeoff between peak memory and runtime.

### 3.2. Results on simulated data

On reads with 0.5% error rate, `synpact` has an accuracy close to `minimap2` for all three genomes while running faster and using less memory (Fig. 3). We observe a similar trend for the other error rates (Suppl. Figs. S2–S5). In the category of resource-light tools, BLEND and `mapquik` are also close in accuracy on human and maize, but on rye, the largest and most repetitive genome [14], their accuracy drops substantially below `synpact` and `minimap2` (BLEND to about 85 %, `mapquik` to about 26 %).

**Figure 3.**
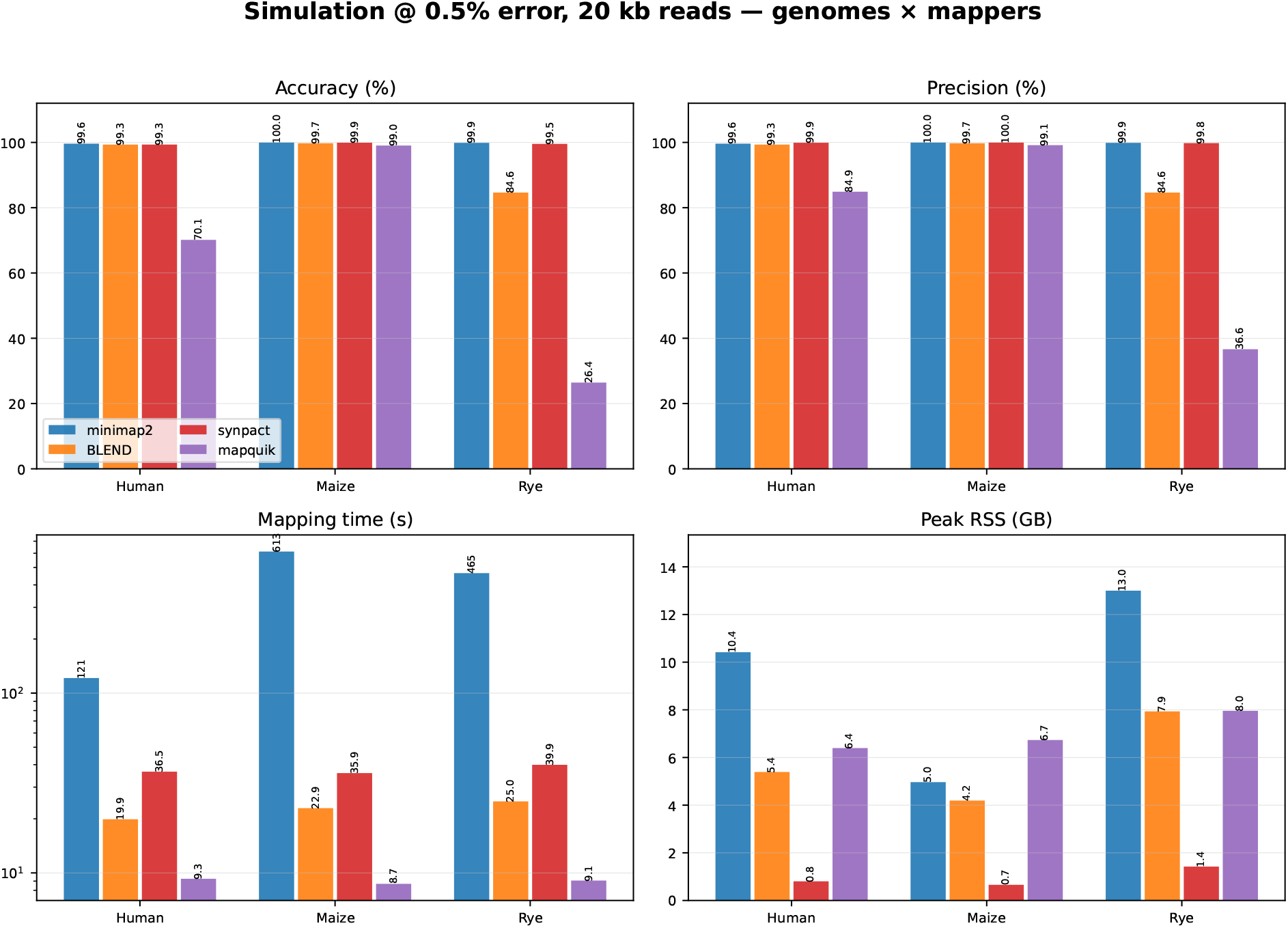
Simulated-read benchmark at 0.5 % error, 20 kb reads. Grouped bars for human, maize and rye, one bar per mapper, across four panels: accuracy, precision, mapping time (log scale) and peak memory.

`synpact` is the only method among the resource-light ones that is accurate on large, repeat-rich genomes. In the mapping-time panel (log scale), `synpact`, BLEND and `mapquik` are all near the fast end, and `minimap2` is much slower. In the peak-memory panel, `synpact` uses substantially less memory than every other mapper for all three genomes. Results for the *Arabidopsis* genome and human chromosome Y in isolation are similar and found in Suppl. Figs. S2–S5. Notably, on chromosome Y (a highly complex repetitive chromosome) `synpact` is almost an order of magnitude faster than BLEND with comparable accuracy. Full numerical results, including wrong-chromosome calls, are found for each reference in Suppl. Tables S1-S5.

### 3.3. Results on real reads

Across the four genomes `synpact` has higher agreement than other resource-light tools, while minimap2 achieves the highest agreement. For example, on rye (Fig. 4), `synpact` reaches 97.3 % agreement, second only to `minimap2` (99.5 %) and far above BLEND (84 %) and `mapquik` (26 %), which place many reads at the wrong locus in this highly repetitive genome, for both the standard HiFi readset and the DeepConsensus readset. On the DeepConsensus reads, `synpact` rises to about 99.6 %, within roughly a third of a percentage point of `minimap2` (99.9 %), while BLEND and `mapquik` stay at about 85 % and 27 %. `synpact` benefits from the corrected reads, because its accuracy depends on the blocks being intact, whereas BLEND and `mapquik` appears to be limited by repeat resolution rather than read quality. `synpact` maps rye reads in about 30 seconds, comparable to BLEND and roughly an order of magnitude faster than `minimap2`. As for the simulated data benchmarks, `synpact` uses far less memory than any of the other tools. For example, on rye it uses about 1.4 GB of memory against `minimap2`’s ∼13 GB and BLEND’s ∼8 GB (Fig. 4).

**Figure 4.**
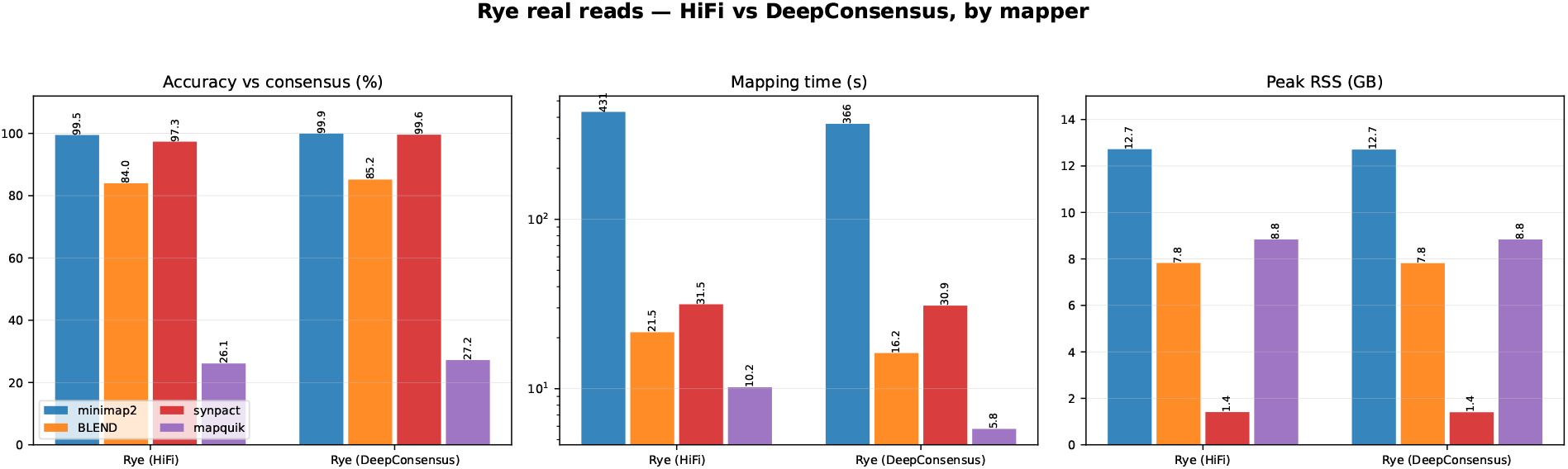
Real rye reads, standard HiFi and DeepConsensus. Accuracy, mapping time and peak memory for both Lo7 readsets, one bar per mapper, scored by consensus of mappers.

Results for the other genomes are shown in Suppl Fig. S6. `synpact` achieves a comparable mapping rate, accuracy, and precision to `minimap2` across genomes, and is more consistent than BLEND and mapquik, which have worse accuracy metrics for the rye genome. `synpact` achieves this accuracy using only a fraction of the memory required by the other tools.

### 3.4. Indexing time and peak memory

Figure 5 shows index build time and peak memory per genome. `synpact`’s indexing time at eight threads is comparable to the other mappers and often faster. It builds the human index in about 35 seconds against 80 seconds for `minimap2` and 58 seconds for BLEND, and is within a few percent of them on rye. However, peak memory during indexing (using eight threads) is one metric that `synpact` does not win. With eight threads it processes several chromosomes in parallel, so its peak build memory is in the same range as `minimap2`, and sometimes higher. This happens where several large chromosomes are processed in parallel. Since `synpact` indexing is fast, we also include `synpact`’s time and peak memory usage during indexing with only one thread. Building with one thread reduces peak memory, e.g., to about 1.7 GB instead of 10.5 GB on human, and 2.0 GB instead of 12.6 GB on maize, at the cost of a longer build time (about 118 seconds instead of 35 on human). The index it produces is identical in both settings. So the peak memory used to build the index can be traded against build time as needed.

**Figure 5.**
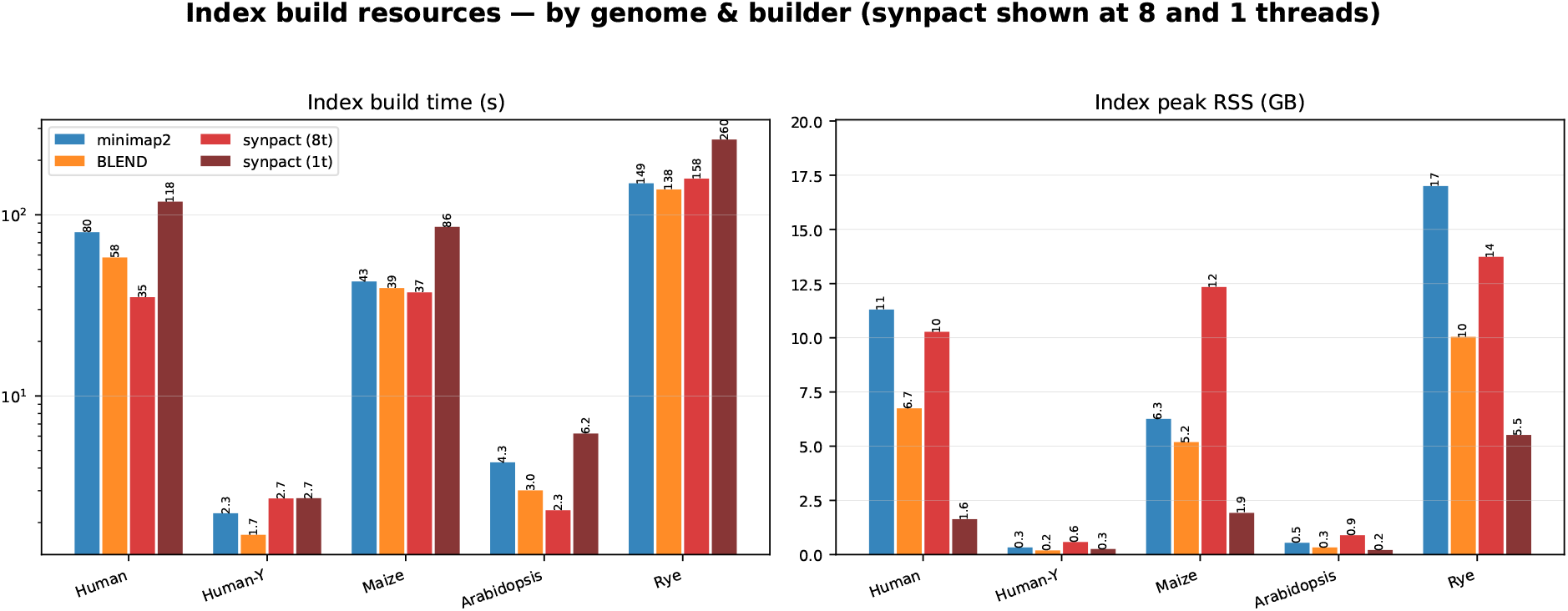
Index build resources by genome and builder. Index build time (log scale) and peak build memory for `minimap2`, BLEND and `synpact`, with `synpact` shown at 8 threads and 1 thread (`synpact1t`). `mapquik` builds its index inline and is not shown.

### 3.5. Mapping to the lungfish genome assembly

The lungfish genome (∼36 Gb) is so large that a default `minimap2` index using `map-hifi` settings does not fit in the available memory and disk, so we ran `minimap2` with sparser seeding (*k*=28, *w*=200) to build an index comparable with `synpact`’s index size. Figure 6 compares accuracy, precision, mapping time, and peak memory of the two tools at 0.5 % error across the four mean read lengths.

**Figure 6.**
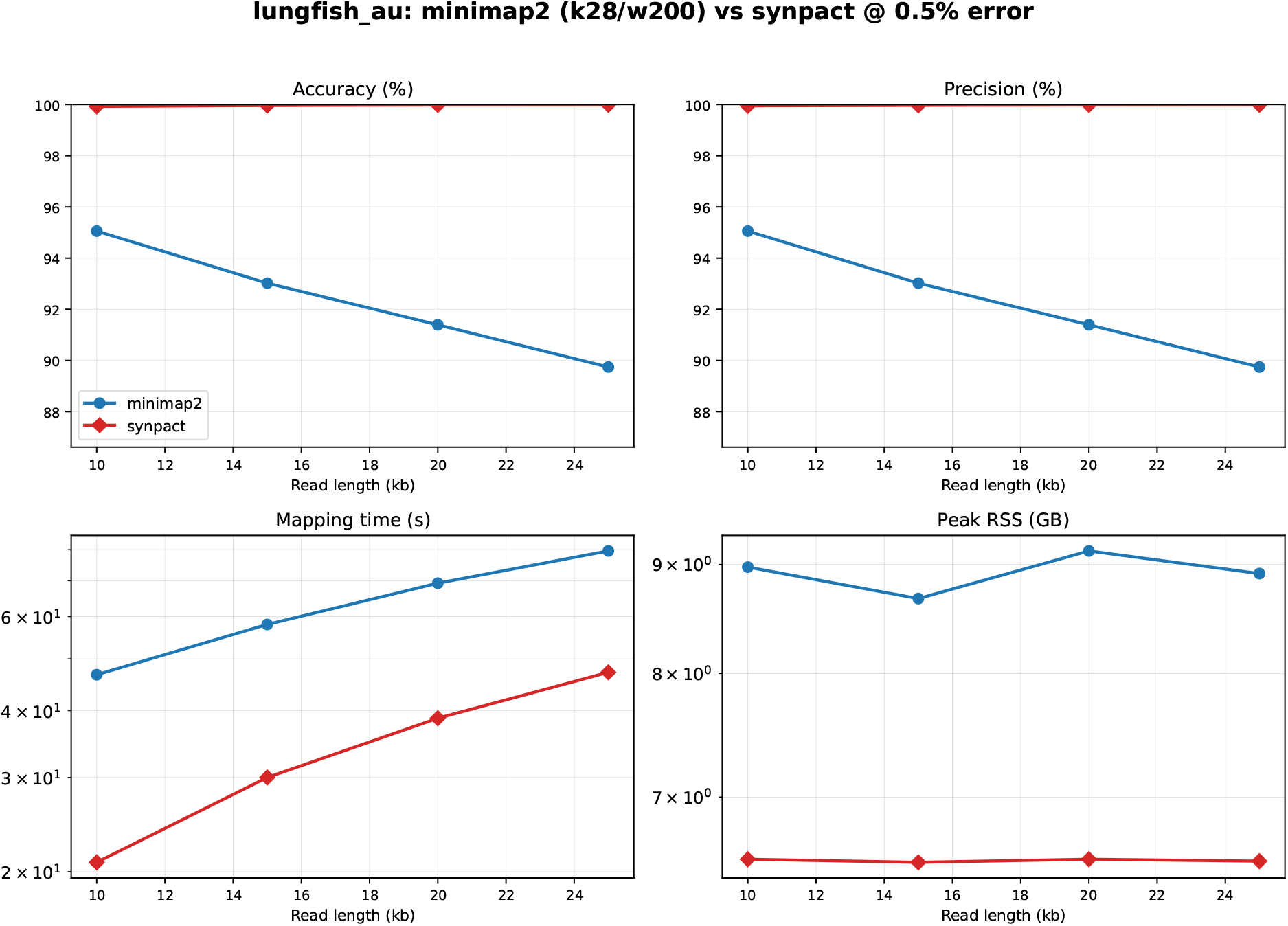
Lungfish benchmark at 0.5 % error. `minimap2` (run with *k*=28, *w*=200) and `synpact` across read length, in four panels: accuracy, precision, mapping time (log scale) and peak memory.

The sparsification of minimap2’s minimizer sampling hurts its accuracy. `synpact` achieves near perfect accuracy with numbers between 99.7–100 % at every read length, whereas the sparsified `minimap2` reaches only about 89–95 %. `synpact` is also cheaper in mapping time and peak memory usage. Similar results were observed for the other error rates (Suppl. Fig. S7).

On the lungfish genome, `synpact` has a slightly slower one-time index build of about 1490 seconds and 14.2 GB of memory, versus about 250 seconds and 12.7 GB for the sparsified `minimap2` index. However, while `synpact` spends more time building its index once, it then maps more accurately, faster, and with less memory.

## 4. Discussion

Inspired by methods using various seed level resolutions for overcoming the specificity-sensitivity tradeoff in read mapping [27], chromosome level mapping [1], and genome assembly [5, 9], we design a seed index that constructs seeds at various resolutions using Locally Consistent Parsing from a syncmer sketch. Our benchmarking results support that for long-read technologies at HiFi error rates, a locally-consistent hierarchical parse of syncmers produces blocks at various resolutions that are efficient at both resolving repeats and being more robust to small levels of errors and variation compared to other lightweight long-read mappers. Indexing only the higher levels in the hierarchy is what gives `synpact` the practical properties of a small index and, thus, low memory usage.

We have demonstrated that `synpact` provides a mapping accuracy comparable with state-of-the-art minimap2 on both simulated (Figs. 3 and 6) and real data (Fig. 4 and Suppl. Fig. S6). Additionally, the index build time is competitive with other methods (Fig. 5). The method is not aimed at improving on `minimap2`’s accuracy, but at reaching comparable accuracy at much lower cost. On small, low-repeat genomes such as *Arabidopsis*, most mappers are already accurate, so `synpact` offers no accuracy advantage. However, the advantage increases with the genome size and repeat content. For example, on rye, `synpact`’s accuracy is close to `minimap2` while the other resource-light mappers, BLEND and `mapquik`, lose substantial accuracy. Moreover, `synpact` acheives this at a fraction of their peak memory. Unlike other lightweight read mappers, `synpact`’s accuracy is relatively stable across reference genomes at various sizes and complexities in our benchmark (Suppl. Fig. S2). The lungfish benchmark demonstrates that `synpact` is also performant on the largest genomes (∼36 Gb) and the maps reads with greater accuracy using less resources than minimap2 with a sparser sketch.

Nevertheless, `synpact`’s method has a clear limit on sequence variability and error rate. Its seed-specificity depends on higher-level blocks being intact, so it is designed for HiFi reads at about 1 % error or below. As the error rate increases, more blocks are corrupted, and `synpact` falls back to finer levels more often. A higher error rate would require storing finer resolution levels, which increases peak memory and decreases speed.

### 4.1. Future work

The output of `synpact` is locus-only (PAF format), which consists of a position and a MAPQ, not a base-level alignment. Reads in ambiguous regions are reported at low MAPQ and could potentially be solved through extension alignment to increase both accuracy and mapping confidence further.

Currently, `synpact` also lacks important long-read features such as supplementary mapping, used frequently, e.g., for structural variation detection. Such features require more sophisticated handling of multiple candidate locations in the diagonal voting method (or considering a different seed clustering approach, such as chaining), and we consider exploring this further.

## 5. Conclusion

We present a HIFi read mapper `synpact`. `synpact` constructs a hierarchical index of seeds at various resolutions (using LCP of syncmers) to find candidate mapping locations. The different seed resolutions make `synpact` both robust to complex repetitive genomes, as well as different error rates within low-error HiFi range (0-1%). `synpact` surpasses the accuracy of other lightweight read mappers, and matches or approaches the accuracy of comparably resource-heavy `minimap2`. `synpact` uses roughly an order of magnitude less memory than the other tools, and also maps reads much faster than minimap2.

## Code availability

`synpact` is accessible on https://github.com/mahmudsami/synpact. All scripts to generate simulations and run benchmarks are available at https://github.com/mahmudsami/hifibenchmark. Accessions for the biological datasets from SRA and ENA are found in Table 1.

## Author contributions

SA conceived the project. SA implemented the idea and performed the benchmarks with input from KS. KS supervised the work. SA and KS wrote the paper.

