## Supplementary file for "synpact: accurate, memory-light PacBio HiFi read mapping via a hierarchy of locally-consistent syncmer blocks"

#### 1 Bounding block length through Deterministic Coin Tossing

We adopt these four rules from LCPan with one exception. A monotone (SSEQ) run can be arbitrarily long, so the rules as stated admit blocks of very uneven length. These long monotone blocks make the per-level distribution of block lengths unbounded. Even though LCPan decreases the bound by doing one step of Deterministic Coin Tossing(DCT) reduction, the bound depends on alphabet size, so it is possible to have 128 blocks (for 64-bit alphabet) in one monotone run. We therefore do not use DCT to label every block, but use it only in long monotone runs, recursively, until we have only 3 possible colors for each block. A strictly monotone run is a proper coloring (neighboring values differ), which DCT reduces, first to at most six colors by the recurrence $c'(i) = 2\pi(i) + b(i)$, where $\pi(i)$ is the lowest bit at which $c(i)$ and $c(i{+}1)$ differ and $b(i)$ is that bit of $c(i)$, then by collapsing the residual colors $\{3, 4, 5\}$ into $\{0, 1, 2\}$ to a proper 3-coloring over $\{0, 1, 2\}$. This 3-coloring is then parsed by the very same local-minimum and local-maximum triplet rules used elsewhere (rules 1–2 above), so the monotone-run blocks are centered triplets identical in form to all other blocks. Because a three-color proper sequence has no two adjacent slope points, these triplets cover the run with blocks of at most three units (Figure S1), so block lengths stay bounded *within* a level. A repetition (RINT) run can be long as well, but we leave it undivided: a run of a single repeated value is not a proper coloring (adjacent values are equal), so the coin-tossing split does not apply, and any positional cut would not be locally consistent; we therefore keep the run as one block. Keeping block sizes uniform across each level allows each level to have a predictable length scale during anchoring and preserves the near-uniform length and spacing that LCP provides at every level.

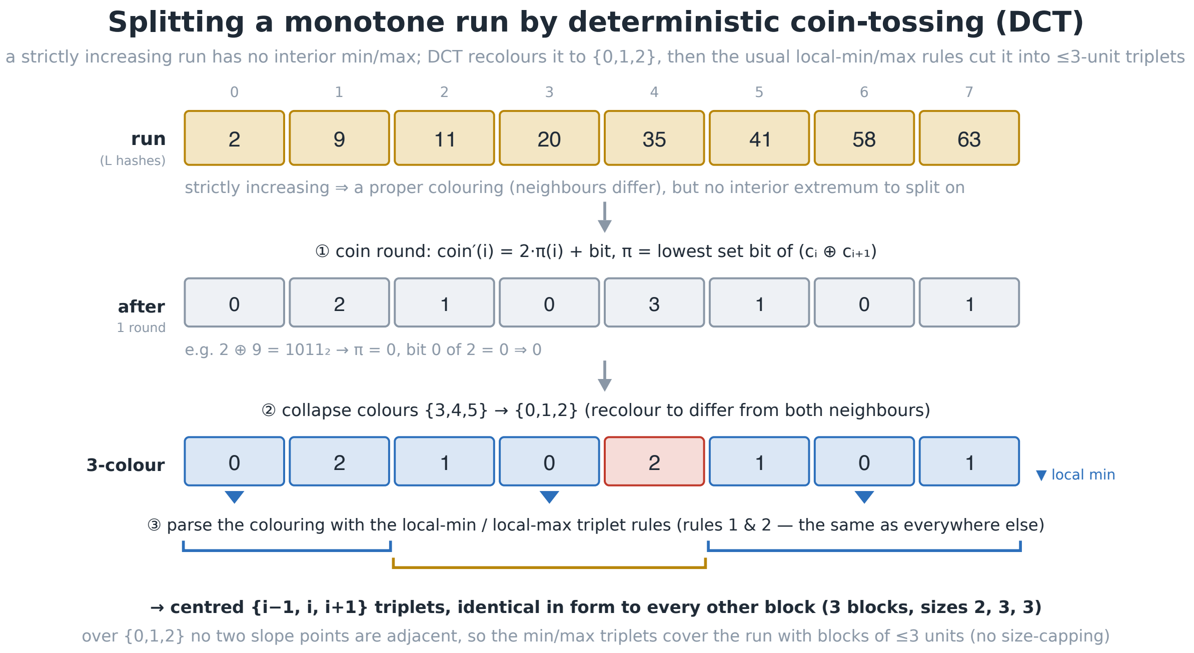


Fig. S1: **Splitting a monotone run by deterministic coin-tossing.** A strictly increasing run has no interior local extremum to split on. One DCT round reduces it via $c'(i) = 2\pi(i) + b(i)$, and collapsing colors $\{3, 4, 5\} \rightarrow \{0, 1, 2\}$ yields a proper 3-coloring. The coloring is then parsed by the usual local-min/local-max triplet rules, giving centered $\{i-1, i, i+1\}$ blocks. Since a three-color sequence has no two adjacent slope points, every block spans $\leq 3$ units without an explicit size cap.

#### 2 Voting algorithm

`synpact` places a read by computing anchor weight along genomic *diagonals*. Each anchor $a = (chr, q, r, w)$ records the read position $q$, the reference position $r$, the chromosome $chr$, and a weight $w$ equal to the block's accumulated syncmer weight. A read coming from a single locus produces anchors that share (approximately) the same diagonal $d(a) = r - q$ (sequencing errors and indels can perturb the diagonal slightly), so a band of width $W$ (default $W = 500$ bp) is used to tolerate smaller variations. Algorithm 1 finds the heaviest such band by a sorted sweep, then re-runs the same sweep over the anchors of every *other* locus to obtain the runner-up weight used for MAPQ.

**Algorithm 1:** VoteLocus — diagonal-voting locus selection

```
Input  : anchors A = {a_i = (chr_i, q_i, r_i, w_i)}; band width W; vote floor τ
Output: best locus (chr*, off*, best) and runner-up weight second
if A = ∅ then return (none, 0)
d(a) ← r_a − q_a                                              // diagonal of an anchor
Sort(A) ascending by the key (chr_a, d(a))
/* Pass 1 -- heaviest diagonal window                                        */
best ← 0;  (lo*, hi*) ← (0,0);  lo ← 0;  sum ← 0
for hi ← 0 to |A| − 1 do
    sum ← sum + w_hi
    while chr_lo ≠ chr_hi or d(a_hi) − d(a_lo) > W do
        sum ← sum − w_lo;  lo ← lo + 1                       // shrink window from the left
    if sum > best then best ← sum; (lo*, hi*) ← (lo, hi)
if best < τ then return (none, 0)
/* Representative offset: anchor of smallest read position in the winning window */
chr* ← chr_hi*;  off* ← d(a_lo*);  q_min ← ∞
for a ∈ A[lo* .. hi*] do
    if q_a < q_min then q_min ← q_a; off* ← d(a)
/* Pass 2 -- heaviest window at a different locus (for MAPQ)                 */
d* ← d(a_hi*)
R ← { a ∈ A : chr_a ≠ chr* or |d(a) − d*| > W }
second ← 0
if R ≠ ∅ then
    lo ← 0;  sum ← 0
    for hi ← 0 to |R| − 1 do
        sum ← sum + w_hi
        while chr_lo ≠ chr_hi or d(R_hi) − d(R_lo) > W do
            sum ← sum − w_lo;  lo ← lo + 1
        second ← max(second, sum)
return ((chr*, off*, best), second)
```

VoteLocus is invoked once per strand. The caller function keeps the higher-scoring strand as the placement and promotes a competing strand's score to *second* only when it sits at a different locus (a different chromosome, or an offset more than $W$ away); a near-identical reverse-complement cluster at the same genomic position is therefore not counted as competition. The final quality is the fraction of the winning weight that is uncontested,

$$\textsc{mapq} \;=\; \min\Big(60,\; \big\lfloor(\mathit{best} - \mathit{second})\cdot 60\,/\,\mathit{best}\big\rfloor\Big),$$

which is 60 for a read with a single dominant locus and falls toward 0 as a second locus approaches the same weight. Sorting dominates the cost, so the procedure runs in $O(n \log n)$ time for $n$ anchors, with both sliding-window passes being linear.

### 3 Figures

For human, human Y chromosome, maize, arabidopsis, and rye simulations, accuracy appears in Figure S2, precision appears in Figure S3, mapping time appears in Figure S4 and peak memory in Figure S5. Results for the lungfish simulation are found in Figure S7. Full results for the real data benchmark are shown in Figure S6.

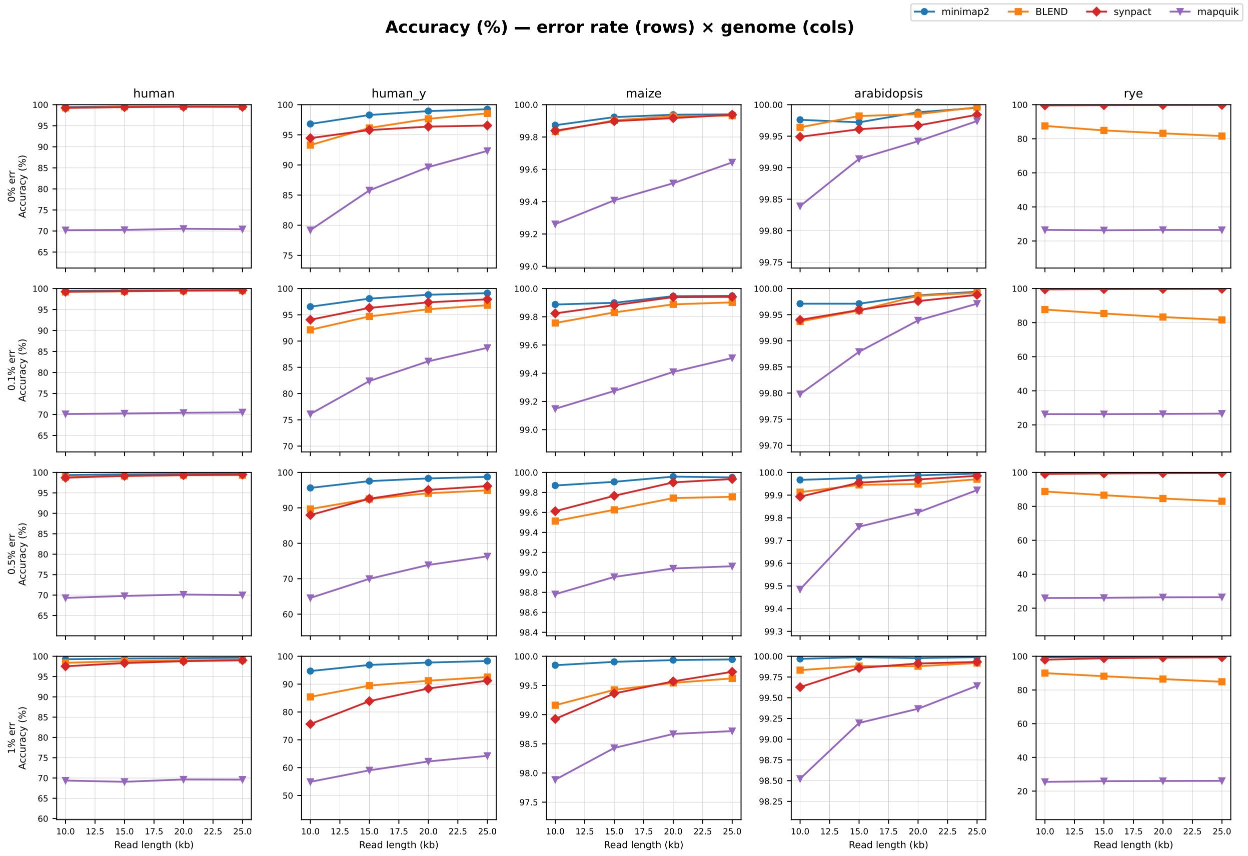


Fig. S2: **Mapping accuracy on simulated reads** Rows are error rate (0, 0.1, 0.5, 1 %), columns are genome (human, human_y, maize, *Arabidopsis*, rye); within each panel the $x$-axis is read length (10–25 kb) and each line is a mapper.

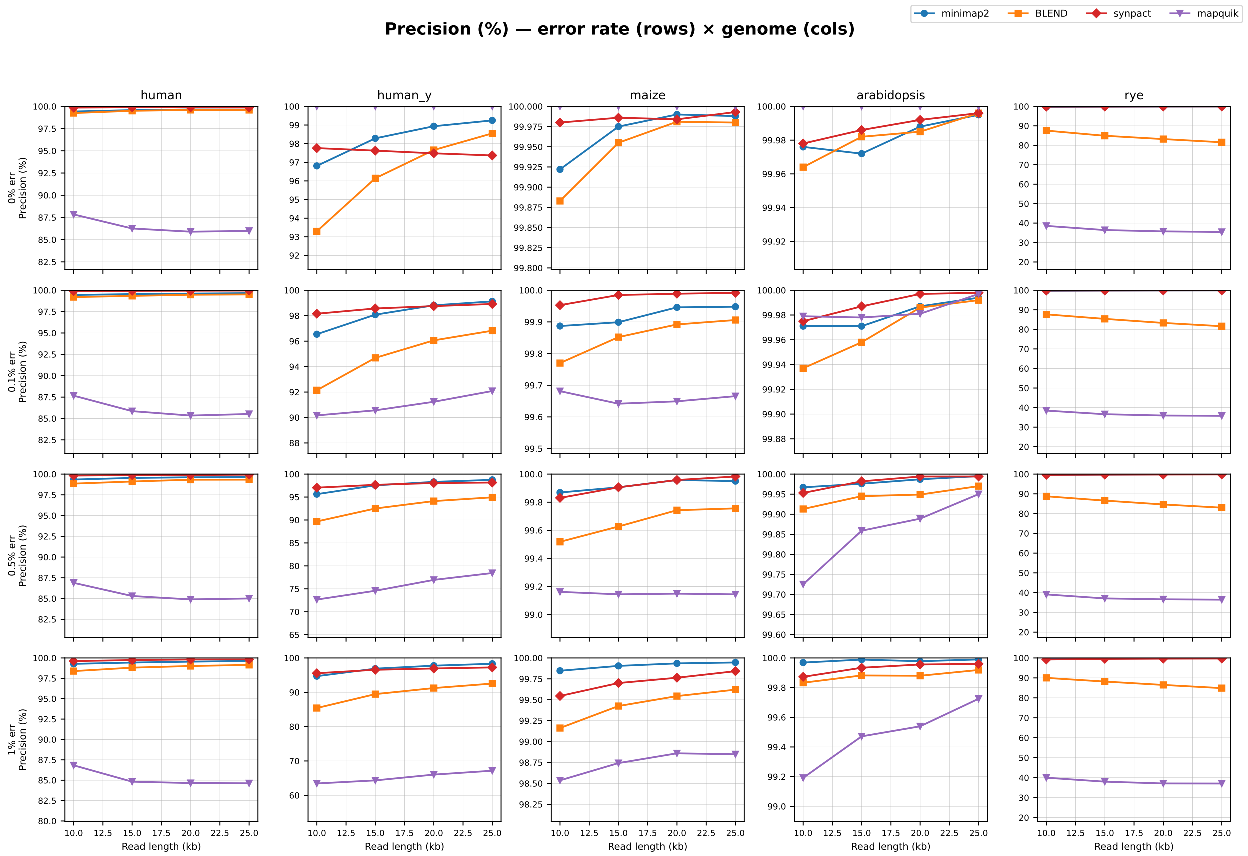


Fig. S3: **Mapping precision on simulated reads** Layout as in Figure S2: rows are error rate, columns are genome, $x$-axis is read length, one line per mapper.

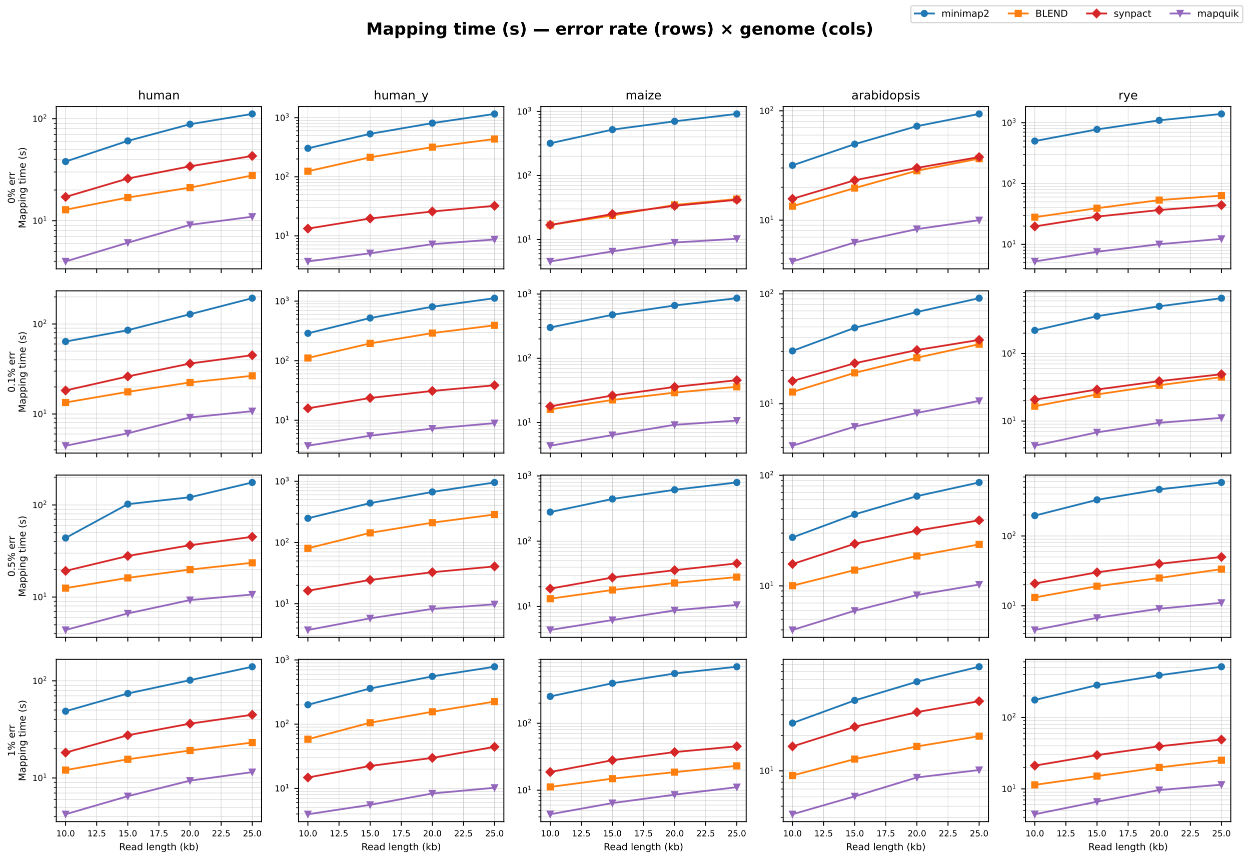


Fig. S4: **Mapping time on simulated reads** (log scale, index loading excluded). Rows are error rate, columns are genome, $x$-axis is read length, one line per mapper.

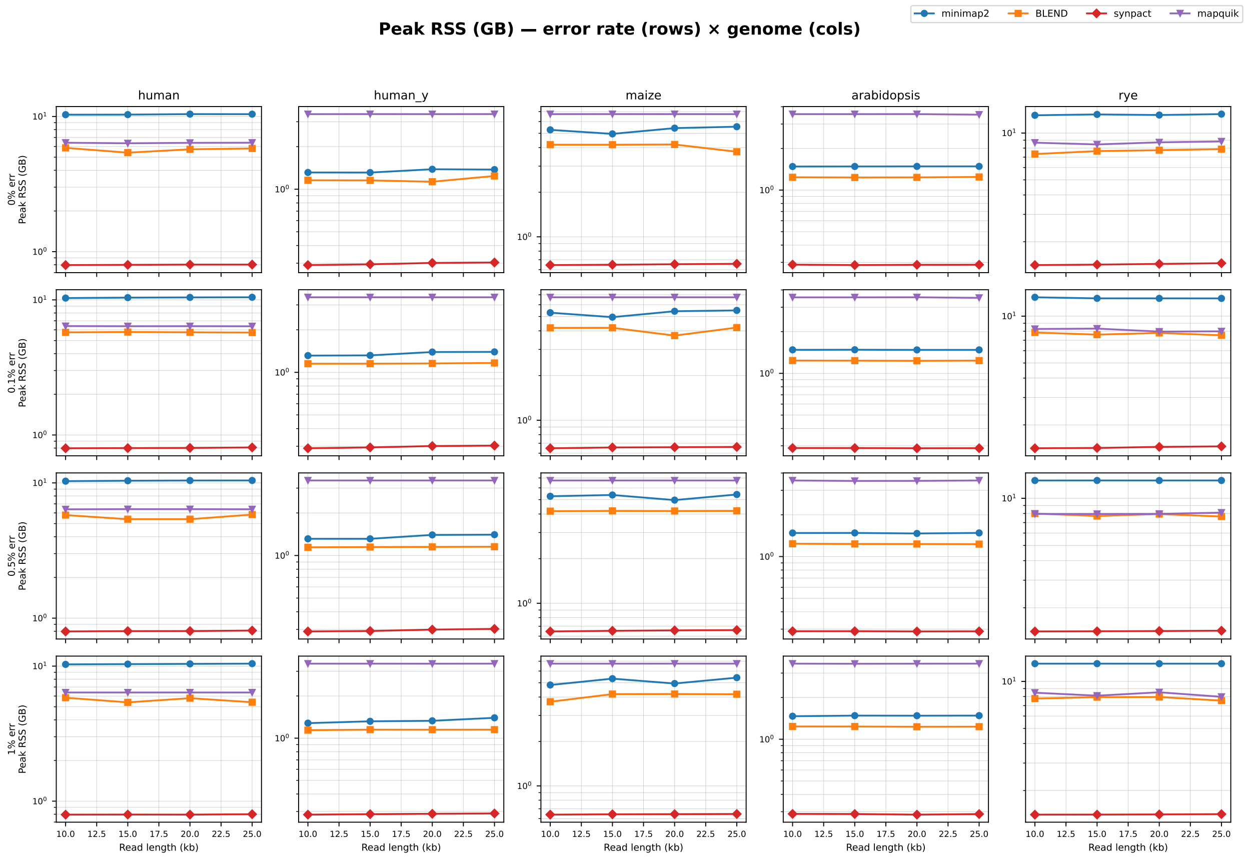


Fig. S5: **Peak mapping memory on simulated reads** (log scale). Rows are error rate, columns are genome, $x$-axis is read length, one line per mapper.

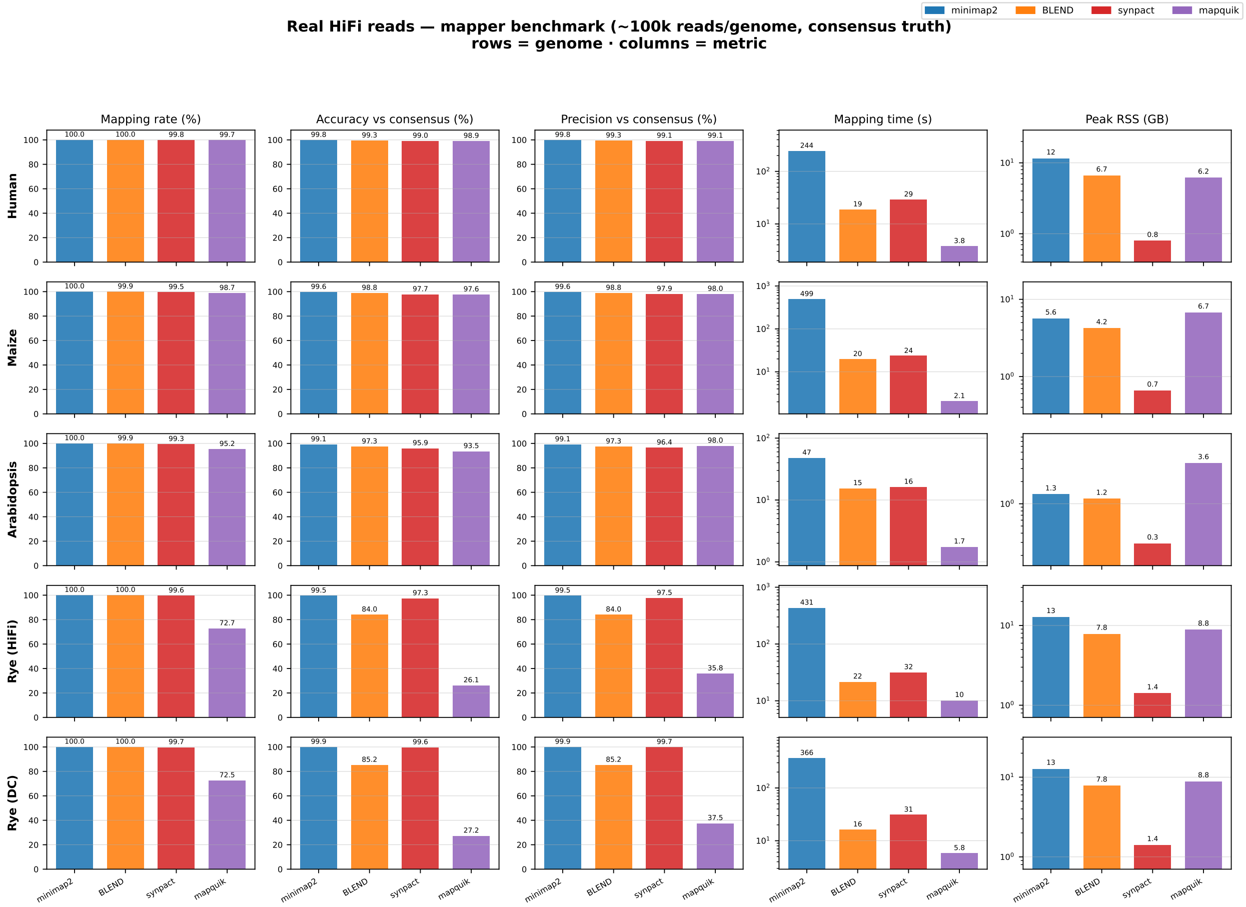


Fig. S6: **Real-data benchmark, all genomes.** Rows are genome / readset (rye split into HiFi and DeepConsensus), columns are metric (accuracy, precision, mapping time, peak memory); each cell is a bar chart with one bar per mapper, scored by consensus of mappers.

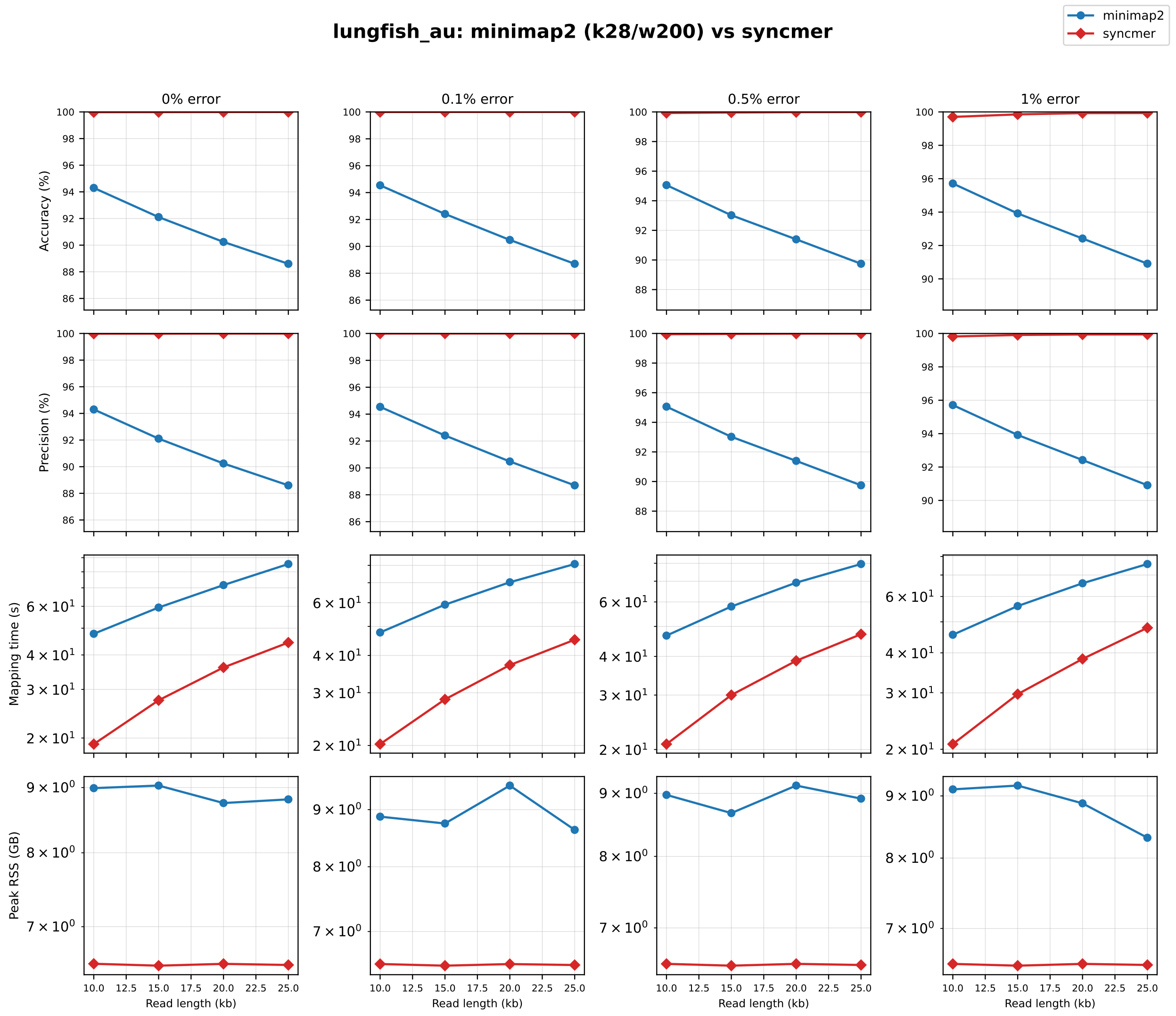


Fig. S7: **Lungfish benchmark, full sweep.** `minimap2` ($k$=28, $w$=200) and `synpact` over all error rates and read lengths: rows are metric (accuracy, precision, mapping time, peak memory), columns are error rate, $x$-axis is read length, one line per mapper.

#### 4 Complete numeric results

Tables S1–S6 report every condition of the benchmark—four error rates × four read lengths × five mappers for each genome, plus the full real-data benchmark—generated directly from the benchmark output.

Table S1: Complete simulated benchmark for *Arabidopsis* (100000 reads per condition). Acc. is over all reads, Prec. over placed reads; W-chr is wrong-chromosome calls; time excludes index loading.

| Mapper | Err. (%) | Len. (kb) | Mapped | Acc. (%) | Prec. (%) | W-chr | Time (s) | RSS (MB) |
|---|---|---|---|---|---|---|---|---|
| minimap2 | 0.0 | 10 | 100000 | 99.98 | 99.98 | 1 | 31.70 | 1514.9 |
| BLEND | 0.0 | 10 | 100000 | 99.96 | 99.96 | 15 | 13.41 | 1266.7 |
| mapquik | 0.0 | 10 | 99839 | 99.84 | 100.00 | 0 | 4.19 | 3622.8 |
| synpact | 0.0 | 10 | 99971 | 99.95 | 99.98 | 6 | 15.68 | 296.0 |
| minimap2 | 0.0 | 15 | 100000 | 99.97 | 99.97 | 0 | 49.55 | 1516.5 |
| BLEND | 0.0 | 15 | 100000 | 99.98 | 99.98 | 0 | 19.65 | 1262.6 |
| mapquik | 0.0 | 15 | 99914 | 99.91 | 100.00 | 0 | 6.25 | 3622.8 |
| synpact | 0.0 | 15 | 99975 | 99.96 | 99.99 | 0 | 23.16 | 293.7 |
| minimap2 | 0.0 | 20 | 100000 | 99.99 | 99.99 | 0 | 72.36 | 1518.0 |
| BLEND | 0.0 | 20 | 100000 | 99.98 | 99.98 | 0 | 28.32 | 1264.6 |
| mapquik | 0.0 | 20 | 99942 | 99.94 | 100.00 | 0 | 8.28 | 3624.8 |
| synpact | 0.0 | 20 | 99975 | 99.97 | 99.99 | 0 | 30.05 | 295.2 |
| minimap2 | 0.0 | 25 | 100000 | 100.00 | 100.00 | 0 | 93.62 | 1518.6 |
| BLEND | 0.0 | 25 | 100000 | 100.00 | 100.00 | 0 | 36.51 | 1272.8 |
| mapquik | 0.0 | 25 | 99974 | 99.97 | 100.00 | 0 | 9.99 | 3590.8 |
| synpact | 0.0 | 25 | 99988 | 99.98 | 100.00 | 0 | 37.74 | 295.5 |
| minimap2 | 0.1 | 10 | 100000 | 99.97 | 99.97 | 3 | 30.25 | 1513.8 |
| BLEND | 0.1 | 10 | 100000 | 99.94 | 99.94 | 32 | 12.78 | 1266.7 |
| mapquik | 0.1 | 10 | 99819 | 99.80 | 99.98 | 2 | 4.15 | 3619.8 |
| synpact | 0.1 | 10 | 99965 | 99.94 | 99.97 | 6 | 16.09 | 296.1 |
| minimap2 | 0.1 | 15 | 100000 | 99.97 | 99.97 | 0 | 49.12 | 1515.9 |
| BLEND | 0.1 | 15 | 100000 | 99.96 | 99.96 | 11 | 19.10 | 1264.6 |
| mapquik | 0.1 | 15 | 99901 | 99.88 | 99.98 | 3 | 6.18 | 3619.9 |
| synpact | 0.1 | 15 | 99972 | 99.96 | 99.99 | 0 | 23.36 | 295.9 |
| minimap2 | 0.1 | 20 | 100000 | 99.99 | 99.99 | 0 | 68.44 | 1512.4 |
| BLEND | 0.1 | 20 | 100000 | 99.99 | 99.99 | 3 | 26.22 | 1261.6 |
| mapquik | 0.1 | 20 | 99958 | 99.94 | 99.98 | 1 | 8.24 | 3624.6 |
| synpact | 0.1 | 20 | 99979 | 99.98 | 100.00 | 0 | 30.82 | 294.9 |
| minimap2 | 0.1 | 25 | 100000 | 99.99 | 99.99 | 0 | 91.35 | 1512.4 |
| BLEND | 0.1 | 25 | 100000 | 99.99 | 99.99 | 4 | 34.82 | 1265.7 |
| mapquik | 0.1 | 25 | 99974 | 99.97 | 100.00 | 0 | 10.60 | 3591.0 |
| synpact | 0.1 | 25 | 99990 | 99.99 | 100.00 | 0 | 38.10 | 295.3 |
| minimap2 | 0.5 | 10 | 100000 | 99.97 | 99.97 | 2 | 27.41 | 1513.9 |
| BLEND | 0.5 | 10 | 100000 | 99.91 | 99.91 | 32 | 10.04 | 1266.7 |
| mapquik | 0.5 | 10 | 99759 | 99.48 | 99.73 | 13 | 3.99 | 3619.6 |
| synpact | 0.5 | 10 | 99940 | 99.89 | 99.95 | 31 | 15.76 | 296.5 |
| minimap2 | 0.5 | 15 | 100000 | 99.98 | 99.98 | 1 | 44.30 | 1515.5 |
| BLEND | 0.5 | 15 | 100000 | 99.94 | 99.94 | 17 | 13.90 | 1262.6 |
| mapquik | 0.5 | 15 | 99902 | 99.76 | 99.86 | 10 | 5.96 | 3590.9 |
| synpact | 0.5 | 15 | 99973 | 99.95 | 99.98 | 6 | 24.02 | 296.6 |
| minimap2 | 0.5 | 20 | 100000 | 99.99 | 99.99 | 0 | 64.77 | 1502.3 |
| BLEND | 0.5 | 20 | 100000 | 99.95 | 99.95 | 8 | 18.64 | 1261.6 |
| mapquik | 0.5 | 20 | 99935 | 99.82 | 99.89 | 8 | 8.26 | 3596.1 |
| synpact | 0.5 | 20 | 99975 | 99.97 | 99.99 | 1 | 31.54 | 295.5 |
| minimap2 | 0.5 | 25 | 100000 | 100.00 | 100.00 | 0 | 86.08 | 1515.9 |
| BLEND | 0.5 | 25 | 100000 | 99.97 | 99.97 | 1 | 23.70 | 1258.5 |
| mapquik | 0.5 | 25 | 99972 | 99.92 | 99.95 | 1 | 10.28 | 3619.8 |

continued on next page

*Table S1 (continued)*

| Mapper | Err. (%) | Len. (kb) | Mapped | Acc. (%) | Prec. (%) | W-chr | Time (s) | RSS (MB) |
|---|---|---|---|---|---|---|---|---|
| synpact | 0.5 | 25 | 99991 | 99.98 | 99.99 | 3 | 39.15 | 296.1 |
| minimap2 | 1.0 | 10 | 100000 | 99.97 | 99.97 | 0 | 25.48 | 1500.7 |
| BLEND | 1.0 | 10 | 100000 | 99.83 | 99.83 | 42 | 9.12 | 1265.7 |
| mapquik | 1.0 | 10 | 99326 | 98.52 | 99.19 | 46 | 4.27 | 3592.8 |
| synpact | 1.0 | 10 | 99755 | 99.63 | 99.87 | 91 | 16.13 | 296.3 |
| minimap2 | 1.0 | 15 | 100000 | 99.99 | 99.99 | 0 | 39.72 | 1515.5 |
| BLEND | 1.0 | 15 | 100000 | 99.88 | 99.88 | 26 | 12.58 | 1264.6 |
| mapquik | 1.0 | 15 | 99723 | 99.20 | 99.47 | 24 | 6.06 | 3585.6 |
| synpact | 1.0 | 15 | 99925 | 99.86 | 99.93 | 44 | 23.69 | 295.8 |
| minimap2 | 1.0 | 20 | 100000 | 99.98 | 99.98 | 0 | 57.29 | 1513.8 |
| BLEND | 1.0 | 20 | 100000 | 99.88 | 99.88 | 17 | 16.12 | 1259.5 |
| mapquik | 1.0 | 20 | 99827 | 99.37 | 99.54 | 31 | 8.78 | 3590.8 |
| synpact | 1.0 | 20 | 99957 | 99.91 | 99.96 | 31 | 31.59 | 292.7 |
| minimap2 | 1.0 | 25 | 100000 | 99.99 | 99.99 | 0 | 76.83 | 1515.5 |
| BLEND | 1.0 | 25 | 100000 | 99.92 | 99.92 | 7 | 19.71 | 1261.6 |
| mapquik | 1.0 | 25 | 99919 | 99.64 | 99.72 | 14 | 10.15 | 3592.8 |
| synpact | 1.0 | 25 | 99972 | 99.93 | 99.96 | 31 | 39.12 | 295.5 |

Table S2: Complete simulated benchmark for Human (100000 reads per condition). Acc. is over all reads, Prec. over placed reads; W-chr is wrong-chromosome calls; time excludes index loading.

| Mapper | Err. (%) | Len. (kb) | Mapped | Acc. (%) | Prec. (%) | W-chr | Time (s) | RSS (MB) |
|---|---|---|---|---|---|---|---|---|
| minimap2 | 0.0 | 10 | 100000 | 99.42 | 99.42 | 8 | 37.99 | 10558.5 |
| BLEND | 0.0 | 10 | 100000 | 99.25 | 99.25 | 25 | 12.81 | 6003.7 |
| mapquik | 0.0 | 10 | 79917 | 70.20 | 87.84 | 9718 | 3.99 | 6547.5 |
| synpact | 0.0 | 10 | 99371 | 99.24 | 99.87 | 2 | 17.10 | 814.7 |
| minimap2 | 0.0 | 15 | 100000 | 99.58 | 99.58 | 0 | 60.58 | 10574.8 |
| BLEND | 0.0 | 15 | 100000 | 99.50 | 99.50 | 8 | 16.86 | 5529.6 |
| mapquik | 0.0 | 15 | 81456 | 70.26 | 86.26 | 11194 | 6.08 | 6487.5 |
| synpact | 0.0 | 15 | 99521 | 99.42 | 99.90 | 2 | 25.91 | 818.2 |
| minimap2 | 0.0 | 20 | 100000 | 99.66 | 99.66 | 0 | 88.23 | 10665.0 |
| BLEND | 0.0 | 20 | 100000 | 99.60 | 99.60 | 4 | 21.11 | 5854.2 |
| mapquik | 0.0 | 20 | 82113 | 70.53 | 85.90 | 11580 | 9.11 | 6535.6 |
| synpact | 0.0 | 20 | 99610 | 99.50 | 99.89 | 1 | 34.13 | 821.6 |
| minimap2 | 0.0 | 25 | 100000 | 99.64 | 99.64 | 0 | 111.33 | 10655.2 |
| BLEND | 0.0 | 25 | 100000 | 99.60 | 99.60 | 1 | 27.75 | 5942.3 |
| mapquik | 0.0 | 25 | 81906 | 70.43 | 85.99 | 11477 | 10.95 | 6549.1 |
| synpact | 0.0 | 25 | 99602 | 99.48 | 99.88 | 0 | 43.11 | 821.8 |
| minimap2 | 0.1 | 10 | 100000 | 99.45 | 99.45 | 7 | 63.88 | 10558.5 |
| BLEND | 0.1 | 10 | 100000 | 99.20 | 99.20 | 29 | 13.43 | 5887.0 |
| mapquik | 0.1 | 10 | 79968 | 70.11 | 87.67 | 9636 | 4.45 | 6549.1 |
| synpact | 0.1 | 10 | 99340 | 99.24 | 99.90 | 3 | 18.27 | 816.4 |
| minimap2 | 0.1 | 15 | 100000 | 99.55 | 99.55 | 1 | 85.34 | 10641.4 |
| BLEND | 0.1 | 15 | 100000 | 99.34 | 99.34 | 15 | 17.63 | 5919.7 |
| mapquik | 0.1 | 15 | 81822 | 70.25 | 85.86 | 11315 | 6.11 | 6535.1 |
| synpact | 0.1 | 15 | 99463 | 99.39 | 99.93 | 0 | 26.15 | 818.8 |
| minimap2 | 0.1 | 20 | 100000 | 99.61 | 99.61 | 1 | 128.44 | 10680.3 |
| BLEND | 0.1 | 20 | 100000 | 99.48 | 99.48 | 3 | 22.41 | 5892.1 |
| mapquik | 0.1 | 20 | 82498 | 70.41 | 85.34 | 11848 | 9.16 | 6539.1 |
| synpact | 0.1 | 20 | 99569 | 99.52 | 99.95 | 1 | 36.39 | 821.6 |
| minimap2 | 0.1 | 25 | 100000 | 99.66 | 99.66 | 0 | 193.53 | 10702.3 |
| BLEND | 0.1 | 25 | 100000 | 99.51 | 99.51 | 7 | 26.58 | 5867.5 |

continued on next page

*Table S2 (continued)*

| Mapper | Err. (%) | Len. (kb) | Mapped | Acc. (%) | Prec. (%) | W-chr | Time (s) | RSS (MB) |
|---|---|---|---|---|---|---|---|---|
| mapquik | 0.1 | 25 | 82446 | 70.51 | 85.52 | 11726 | 10.75 | 6529.4 |
| synpact | 0.1 | 25 | 99624 | 99.59 | 99.96 | 2 | 45.01 | 828.1 |
| minimap2 | 0.5 | 10 | 100000 | 99.34 | 99.34 | 5 | 43.85 | 10550.3 |
| BLEND | 0.5 | 10 | 100000 | 98.85 | 98.85 | 39 | 12.50 | 5921.8 |
| mapquik | 0.5 | 10 | 79769 | 69.31 | 86.89 | 9718 | 4.39 | 6531.1 |
| synpact | 0.5 | 10 | 98883 | 98.71 | 99.82 | 5 | 19.24 | 816.4 |
| minimap2 | 0.5 | 15 | 100000 | 99.54 | 99.54 | 8 | 102.10 | 10628.1 |
| BLEND | 0.5 | 15 | 100000 | 99.11 | 99.11 | 31 | 16.16 | 5520.4 |
| mapquik | 0.5 | 15 | 81816 | 69.79 | 85.30 | 11369 | 6.64 | 6548.8 |
| synpact | 0.5 | 15 | 99277 | 99.16 | 99.88 | 3 | 27.93 | 821.1 |
| minimap2 | 0.5 | 20 | 100000 | 99.62 | 99.62 | 0 | 121.27 | 10667.3 |
| BLEND | 0.5 | 20 | 100000 | 99.32 | 99.32 | 18 | 19.87 | 5520.4 |
| mapquik | 0.5 | 20 | 82618 | 70.14 | 84.89 | 11823 | 9.27 | 6547.1 |
| synpact | 0.5 | 20 | 99451 | 99.34 | 99.89 | 2 | 36.55 | 822.0 |
| minimap2 | 0.5 | 25 | 100000 | 99.63 | 99.63 | 1 | 175.53 | 10673.2 |
| BLEND | 0.5 | 25 | 100000 | 99.33 | 99.33 | 22 | 23.57 | 5972.0 |
| mapquik | 0.5 | 25 | 82328 | 69.99 | 85.01 | 11711 | 10.66 | 6537.1 |
| synpact | 0.5 | 25 | 99549 | 99.48 | 99.93 | 3 | 45.10 | 829.2 |
| minimap2 | 1.0 | 10 | 100000 | 99.28 | 99.28 | 9 | 48.63 | 10549.2 |
| BLEND | 1.0 | 10 | 100000 | 98.39 | 98.39 | 70 | 12.06 | 5977.1 |
| mapquik | 1.0 | 10 | 79890 | 69.37 | 86.83 | 9633 | 4.23 | 6533.4 |
| synpact | 1.0 | 10 | 97909 | 97.53 | 99.62 | 27 | 18.25 | 813.6 |
| minimap2 | 1.0 | 15 | 100000 | 99.44 | 99.44 | 5 | 74.06 | 10597.4 |
| BLEND | 1.0 | 15 | 100000 | 98.81 | 98.81 | 44 | 15.57 | 5519.4 |
| mapquik | 1.0 | 15 | 81426 | 69.07 | 84.82 | 11449 | 6.49 | 6533.1 |
| synpact | 1.0 | 15 | 98590 | 98.34 | 99.74 | 19 | 27.51 | 814.8 |
| minimap2 | 1.0 | 20 | 100000 | 99.55 | 99.55 | 1 | 101.56 | 10640.6 |
| BLEND | 1.0 | 20 | 100000 | 99.01 | 99.01 | 29 | 19.18 | 5929.0 |
| mapquik | 1.0 | 20 | 82261 | 69.64 | 84.65 | 11718 | 9.39 | 6533.6 |
| synpact | 1.0 | 20 | 98973 | 98.79 | 99.82 | 8 | 36.23 | 813.8 |
| minimap2 | 1.0 | 25 | 100000 | 99.64 | 99.64 | 0 | 139.62 | 10680.3 |
| BLEND | 1.0 | 25 | 100000 | 99.15 | 99.15 | 32 | 23.15 | 5532.7 |
| mapquik | 1.0 | 25 | 82247 | 69.60 | 84.62 | 11774 | 11.48 | 6531.1 |
| synpact | 1.0 | 25 | 99189 | 99.02 | 99.83 | 6 | 44.73 | 819.7 |

Table S3: Complete simulated benchmark for Human (chrY) (100000 reads per condition). Acc. is over all reads, Prec. over placed reads; W-chr is wrong-chromosome calls; time excludes index loading.

| Mapper | Err. (%) | Len. (kb) | Mapped | Acc. (%) | Prec. (%) | W-chr | Time (s) | RSS (MB) |
|---|---|---|---|---|---|---|---|---|
| minimap2 | 0.0 | 10 | 100000 | 96.81 | 96.81 | 0 | 303.17 | 1345.5 |
| BLEND | 0.0 | 10 | 100000 | 93.30 | 93.30 | 0 | 123.98 | 1184.8 |
| mapquik | 0.0 | 10 | 79191 | 79.19 | 100.00 | 0 | 3.69 | 3477.6 |
| synpact | 0.0 | 10 | 96613 | 94.45 | 97.76 | 0 | 13.26 | 297.8 |
| minimap2 | 0.0 | 15 | 100000 | 98.28 | 98.28 | 0 | 532.28 | 1343.5 |
| BLEND | 0.0 | 15 | 100000 | 96.14 | 96.14 | 0 | 213.21 | 1182.7 |
| mapquik | 0.0 | 15 | 85782 | 85.78 | 100.00 | 0 | 5.07 | 3479.6 |
| synpact | 0.0 | 15 | 98119 | 95.79 | 97.63 | 0 | 19.63 | 301.5 |
| minimap2 | 0.0 | 20 | 100000 | 98.94 | 98.94 | 0 | 809.39 | 1417.7 |
| BLEND | 0.0 | 20 | 100000 | 97.66 | 97.66 | 0 | 317.33 | 1157.1 |
| mapquik | 0.0 | 20 | 89648 | 89.65 | 100.00 | 0 | 7.23 | 3477.7 |
| synpact | 0.0 | 20 | 98851 | 96.36 | 97.48 | 0 | 25.83 | 308.5 |
| minimap2 | 0.0 | 25 | 100000 | 99.25 | 99.25 | 0 | 1157.97 | 1410.0 |

continued on next page

*Table S3 (continued)*

| Mapper | Err. (%) | Len. (kb) | Mapped | Acc. (%) | Prec. (%) | W-chr | Time (s) | RSS (MB) |
|---|---|---|---|---|---|---|---|---|
| BLEND | 0.0 | 25 | 100000 | 98.55 | 98.55 | 0 | 435.18 | 1270.8 |
| mapquik | 0.0 | 25 | 92330 | 92.33 | 100.00 | 0 | 8.68 | 3479.7 |
| synpact | 0.0 | 25 | 99152 | 96.54 | 97.37 | 0 | 32.24 | 310.7 |
| minimap2 | 0.1 | 10 | 100000 | 96.55 | 96.55 | 0 | 287.85 | 1345.5 |
| BLEND | 0.1 | 10 | 100000 | 92.14 | 92.14 | 0 | 110.89 | 1178.6 |
| mapquik | 0.1 | 10 | 84409 | 76.11 | 90.17 | 0 | 3.73 | 3477.6 |
| synpact | 0.1 | 10 | 95828 | 94.06 | 98.16 | 0 | 15.86 | 297.3 |
| minimap2 | 0.1 | 15 | 100000 | 98.08 | 98.08 | 0 | 519.72 | 1349.8 |
| BLEND | 0.1 | 15 | 100000 | 94.69 | 94.69 | 0 | 195.04 | 1179.6 |
| mapquik | 0.1 | 15 | 90977 | 82.39 | 90.56 | 0 | 5.50 | 3477.7 |
| synpact | 0.1 | 15 | 97719 | 96.32 | 98.57 | 0 | 23.63 | 301.5 |
| minimap2 | 0.1 | 20 | 100000 | 98.82 | 98.82 | 0 | 804.62 | 1425.4 |
| BLEND | 0.1 | 20 | 100000 | 96.06 | 96.06 | 0 | 290.50 | 1183.7 |
| mapquik | 0.1 | 20 | 94421 | 86.14 | 91.23 | 0 | 7.23 | 3477.6 |
| synpact | 0.1 | 20 | 98601 | 97.37 | 98.75 | 0 | 31.04 | 308.3 |
| minimap2 | 0.1 | 25 | 100000 | 99.12 | 99.12 | 0 | 1119.36 | 1428.8 |
| BLEND | 0.1 | 25 | 100000 | 96.82 | 96.82 | 0 | 392.27 | 1193.0 |
| mapquik | 0.1 | 25 | 96341 | 88.71 | 92.08 | 0 | 8.93 | 3477.6 |
| synpact | 0.1 | 25 | 99018 | 97.95 | 98.92 | 0 | 38.61 | 310.6 |
| minimap2 | 0.5 | 10 | 100000 | 95.63 | 95.63 | 0 | 248.40 | 1344.8 |
| BLEND | 0.5 | 10 | 100000 | 89.68 | 89.68 | 0 | 80.25 | 1170.4 |
| mapquik | 0.5 | 10 | 88870 | 64.58 | 72.66 | 0 | 3.71 | 3477.7 |
| synpact | 0.5 | 10 | 90646 | 87.97 | 97.04 | 0 | 16.28 | 297.2 |
| minimap2 | 0.5 | 15 | 100000 | 97.55 | 97.55 | 0 | 439.86 | 1344.6 |
| BLEND | 0.5 | 15 | 100000 | 92.50 | 92.50 | 0 | 143.62 | 1175.6 |
| mapquik | 0.5 | 15 | 93818 | 69.97 | 74.58 | 0 | 5.76 | 3477.6 |
| synpact | 0.5 | 15 | 94795 | 92.58 | 97.66 | 0 | 24.41 | 299.4 |
| minimap2 | 0.5 | 20 | 100000 | 98.31 | 98.31 | 0 | 669.96 | 1432.6 |
| BLEND | 0.5 | 20 | 100000 | 94.12 | 94.12 | 0 | 210.75 | 1178.6 |
| mapquik | 0.5 | 20 | 96017 | 73.88 | 76.95 | 0 | 8.21 | 3477.6 |
| synpact | 0.5 | 20 | 96973 | 95.08 | 98.05 | 0 | 32.66 | 306.3 |
| minimap2 | 0.5 | 25 | 100000 | 98.75 | 98.75 | 0 | 956.64 | 1438.1 |
| BLEND | 0.5 | 25 | 100000 | 94.94 | 94.94 | 0 | 286.43 | 1181.7 |
| mapquik | 0.5 | 25 | 97279 | 76.31 | 78.44 | 0 | 9.76 | 3477.7 |
| synpact | 0.5 | 25 | 97944 | 96.14 | 98.16 | 0 | 40.63 | 309.9 |
| minimap2 | 1.0 | 10 | 100000 | 94.69 | 94.69 | 0 | 200.67 | 1310.7 |
| BLEND | 1.0 | 10 | 100000 | 85.42 | 85.42 | 0 | 58.27 | 1168.4 |
| mapquik | 1.0 | 10 | 86533 | 54.90 | 63.45 | 0 | 3.96 | 3479.7 |
| synpact | 1.0 | 10 | 79169 | 75.65 | 95.56 | 0 | 14.76 | 291.8 |
| minimap2 | 1.0 | 15 | 100000 | 96.89 | 96.89 | 0 | 359.37 | 1350.7 |
| BLEND | 1.0 | 15 | 100000 | 89.46 | 89.46 | 0 | 105.35 | 1177.6 |
| mapquik | 1.0 | 15 | 91754 | 59.04 | 64.34 | 0 | 5.55 | 3479.5 |
| synpact | 1.0 | 15 | 86875 | 83.88 | 96.55 | 0 | 22.36 | 294.1 |
| minimap2 | 1.0 | 20 | 100000 | 97.75 | 97.75 | 0 | 554.30 | 1361.9 |
| BLEND | 1.0 | 20 | 100000 | 91.23 | 91.23 | 0 | 156.00 | 1176.6 |
| mapquik | 1.0 | 20 | 94238 | 62.22 | 66.03 | 0 | 8.34 | 3477.7 |
| synpact | 1.0 | 20 | 91224 | 88.43 | 96.94 | 0 | 29.84 | 295.9 |
| minimap2 | 1.0 | 25 | 100000 | 98.31 | 98.31 | 0 | 783.86 | 1430.5 |
| BLEND | 1.0 | 25 | 100000 | 92.54 | 92.54 | 0 | 224.91 | 1177.6 |
| mapquik | 1.0 | 25 | 95660 | 64.24 | 67.16 | 0 | 10.23 | 3480.6 |
| synpact | 1.0 | 25 | 93847 | 91.26 | 97.24 | 0 | 44.37 | 297.5 |

Table S4: Complete simulated benchmark for Maize (100000 reads per condition). Acc. is over all reads, Prec. over placed reads; W-chr is wrong-chromosome calls; time excludes index loading.

| Mapper | Err. (%) | Len. (kb) | Mapped | Acc. (%) | Prec. (%) | W-chr | Time (s) | RSS (MB) |
|---|---|---|---|---|---|---|---|---|
| minimap2 | 0.0 | 10 | 99951 | 99.87 | 99.92 | 0 | 317.86 | 5400.7 |
| BLEND | 0.0 | 10 | 99951 | 99.83 | 99.88 | 2 | 16.85 | 4287.5 |
| mapquik | 0.0 | 10 | 99261 | 99.26 | 100.00 | 0 | 4.55 | 6897.0 |
| synpact | 0.0 | 10 | 99859 | 99.84 | 99.98 | 0 | 16.88 | 659.7 |
| minimap2 | 0.0 | 15 | 99948 | 99.92 | 99.97 | 0 | 519.46 | 5072.3 |
| BLEND | 0.0 | 15 | 99948 | 99.90 | 99.95 | 1 | 23.72 | 4286.5 |
| mapquik | 0.0 | 15 | 99408 | 99.41 | 100.00 | 0 | 6.52 | 6898.9 |
| synpact | 0.0 | 15 | 99912 | 99.90 | 99.99 | 0 | 25.00 | 663.6 |
| minimap2 | 0.0 | 20 | 99948 | 99.94 | 99.99 | 0 | 700.27 | 5547.1 |
| BLEND | 0.0 | 20 | 99948 | 99.93 | 99.98 | 0 | 34.67 | 4301.8 |
| mapquik | 0.0 | 20 | 99514 | 99.51 | 100.00 | 0 | 8.98 | 6896.9 |
| synpact | 0.0 | 20 | 99933 | 99.92 | 99.98 | 0 | 33.59 | 668.6 |
| minimap2 | 0.0 | 25 | 99952 | 99.94 | 99.99 | 0 | 913.50 | 5671.3 |
| BLEND | 0.0 | 25 | 99952 | 99.93 | 99.98 | 0 | 42.81 | 3842.0 |
| mapquik | 0.0 | 25 | 99643 | 99.64 | 100.00 | 0 | 10.27 | 6894.8 |
| synpact | 0.0 | 25 | 99945 | 99.94 | 99.99 | 0 | 41.68 | 672.7 |
| minimap2 | 0.1 | 10 | 100000 | 99.89 | 99.89 | 25 | 301.62 | 5430.3 |
| BLEND | 0.1 | 10 | 99986 | 99.76 | 99.77 | 21 | 16.03 | 4290.6 |
| mapquik | 0.1 | 10 | 99465 | 99.15 | 99.68 | 40 | 4.36 | 6894.9 |
| synpact | 0.1 | 10 | 99871 | 99.82 | 99.95 | 2 | 17.88 | 663.2 |
| minimap2 | 0.1 | 15 | 100000 | 99.90 | 99.90 | 37 | 475.95 | 5066.9 |
| BLEND | 0.1 | 15 | 99979 | 99.83 | 99.85 | 23 | 22.46 | 4294.7 |
| mapquik | 0.1 | 15 | 99631 | 99.27 | 99.64 | 39 | 6.39 | 6894.9 |
| synpact | 0.1 | 15 | 99897 | 99.88 | 99.98 | 0 | 26.36 | 671.6 |
| minimap2 | 0.1 | 20 | 100000 | 99.95 | 99.95 | 20 | 663.87 | 5553.2 |
| BLEND | 0.1 | 20 | 99996 | 99.89 | 99.89 | 22 | 29.20 | 3807.2 |
| mapquik | 0.1 | 20 | 99759 | 99.41 | 99.65 | 17 | 9.22 | 6894.8 |
| synpact | 0.1 | 20 | 99950 | 99.94 | 99.99 | 0 | 35.86 | 674.0 |
| minimap2 | 0.1 | 25 | 100000 | 99.95 | 99.95 | 28 | 861.76 | 5625.2 |
| BLEND | 0.1 | 25 | 99996 | 99.90 | 99.91 | 26 | 36.01 | 4311.0 |
| mapquik | 0.1 | 25 | 99842 | 99.51 | 99.67 | 17 | 10.71 | 6894.9 |
| synpact | 0.1 | 25 | 99949 | 99.94 | 99.99 | 0 | 45.63 | 677.0 |
| minimap2 | 0.5 | 10 | 100000 | 99.87 | 99.87 | 27 | 279.07 | 5396.9 |
| BLEND | 0.5 | 10 | 99995 | 99.51 | 99.52 | 30 | 13.14 | 4280.3 |
| mapquik | 0.5 | 10 | 99616 | 98.78 | 99.16 | 92 | 4.35 | 6895.0 |
| synpact | 0.5 | 10 | 99781 | 99.61 | 99.83 | 11 | 18.73 | 660.9 |
| minimap2 | 0.5 | 15 | 100000 | 99.91 | 99.91 | 40 | 443.31 | 5504.0 |
| BLEND | 0.5 | 15 | 99999 | 99.63 | 99.63 | 41 | 17.86 | 4297.7 |
| mapquik | 0.5 | 15 | 99808 | 98.95 | 99.14 | 43 | 6.18 | 6896.9 |
| synpact | 0.5 | 15 | 99860 | 99.77 | 99.91 | 0 | 27.71 | 667.6 |
| minimap2 | 0.5 | 20 | 100000 | 99.96 | 99.96 | 18 | 613.20 | 5084.7 |
| BLEND | 0.5 | 20 | 100000 | 99.74 | 99.74 | 21 | 22.91 | 4291.6 |
| mapquik | 0.5 | 20 | 99889 | 99.04 | 99.15 | 31 | 8.69 | 6894.9 |
| synpact | 0.5 | 20 | 99942 | 99.90 | 99.96 | 0 | 35.90 | 672.0 |
| minimap2 | 0.5 | 25 | 100000 | 99.95 | 99.95 | 25 | 793.68 | 5547.2 |
| BLEND | 0.5 | 25 | 100000 | 99.76 | 99.76 | 22 | 28.16 | 4300.8 |
| mapquik | 0.5 | 25 | 99916 | 99.06 | 99.14 | 12 | 10.55 | 6896.9 |
| synpact | 0.5 | 25 | 99951 | 99.93 | 99.98 | 0 | 45.64 | 675.1 |
| minimap2 | 1.0 | 10 | 100000 | 99.85 | 99.85 | 33 | 251.68 | 4943.9 |
| BLEND | 1.0 | 10 | 99999 | 99.16 | 99.16 | 55 | 11.19 | 3806.2 |
| mapquik | 1.0 | 10 | 99339 | 97.89 | 98.54 | 276 | 4.37 | 6894.9 |
| synpact | 1.0 | 10 | 99378 | 98.93 | 99.55 | 103 | 18.69 | 653.8 |

continued on next page

*Table S4 (continued)*

| Mapper | Err. (%) | Len. (kb) | Mapped | Acc. (%) | Prec. (%) | W-chr | Time (s) | RSS (MB) |
|---|---|---|---|---|---|---|---|---|
| minimap2 | 1.0 | 15 | 100000 | 99.91 | 99.91 | 28 | 396.04 | 5457.3 |
| BLEND | 1.0 | 15 | 100000 | 99.42 | 99.42 | 34 | 14.87 | 4288.5 |
| mapquik | 1.0 | 15 | 99682 | 98.43 | 98.74 | 102 | 6.42 | 6894.9 |
| synpact | 1.0 | 15 | 99662 | 99.36 | 99.70 | 12 | 28.04 | 657.3 |
| minimap2 | 1.0 | 20 | 100000 | 99.94 | 99.94 | 25 | 554.18 | 5058.8 |
| BLEND | 1.0 | 20 | 100000 | 99.54 | 99.54 | 24 | 18.61 | 4293.6 |
| mapquik | 1.0 | 20 | 99807 | 98.67 | 98.86 | 33 | 8.58 | 6896.9 |
| synpact | 1.0 | 20 | 99806 | 99.57 | 99.76 | 4 | 37.12 | 658.8 |
| minimap2 | 1.0 | 25 | 100000 | 99.95 | 99.95 | 22 | 699.29 | 5541.9 |
| BLEND | 1.0 | 25 | 100000 | 99.62 | 99.62 | 30 | 23.09 | 4280.3 |
| mapquik | 1.0 | 25 | 99868 | 98.72 | 98.85 | 21 | 11.10 | 6896.9 |
| synpact | 1.0 | 25 | 99892 | 99.73 | 99.84 | 3 | 45.28 | 660.5 |

Table S5: Complete simulated benchmark for Rye (100000 reads per condition). Acc. is over all reads, Prec. over placed reads; W-chr is wrong-chromosome calls; time excludes index loading.

| Mapper | Err. (%) | Len. (kb) | Mapped | Acc. (%) | Prec. (%) | W-chr | Time (s) | RSS (MB) |
|---|---|---|---|---|---|---|---|---|
| minimap2 | 0.0 | 10 | 100000 | 99.74 | 99.74 | 101 | 499.72 | 13320.5 |
| BLEND | 0.0 | 10 | 100000 | 87.55 | 87.55 | 12349 | 27.94 | 7503.9 |
| mapquik | 0.0 | 10 | 68748 | 26.50 | 38.55 | 42244 | 5.24 | 8877.9 |
| synpact | 0.0 | 10 | 99690 | 99.58 | 99.89 | 48 | 19.72 | 1449.7 |
| minimap2 | 0.0 | 15 | 100000 | 99.85 | 99.85 | 60 | 776.08 | 13492.2 |
| BLEND | 0.0 | 15 | 100000 | 84.89 | 84.89 | 15068 | 39.31 | 7837.7 |
| mapquik | 0.0 | 15 | 72195 | 26.29 | 36.41 | 45906 | 7.55 | 8650.2 |
| synpact | 0.0 | 15 | 99799 | 99.71 | 99.91 | 48 | 28.59 | 1460.2 |
| minimap2 | 0.0 | 20 | 100000 | 99.91 | 99.91 | 22 | 1094.26 | 13371.4 |
| BLEND | 0.0 | 20 | 100000 | 83.19 | 83.19 | 16775 | 53.42 | 7947.3 |
| mapquik | 0.0 | 20 | 74129 | 26.49 | 35.73 | 47640 | 10.05 | 8923.1 |
| synpact | 0.0 | 20 | 99859 | 99.81 | 99.95 | 22 | 36.66 | 1475.1 |
| minimap2 | 0.0 | 25 | 100000 | 99.94 | 99.94 | 10 | 1395.00 | 13556.8 |
| BLEND | 0.0 | 25 | 100000 | 81.56 | 81.56 | 18417 | 63.33 | 8071.2 |
| mapquik | 0.0 | 25 | 74675 | 26.46 | 35.44 | 48212 | 12.31 | 9043.0 |
| synpact | 0.0 | 25 | 99893 | 99.85 | 99.96 | 18 | 44.13 | 1490.7 |
| minimap2 | 0.1 | 10 | 100000 | 99.72 | 99.72 | 118 | 219.70 | 13536.3 |
| BLEND | 0.1 | 10 | 100000 | 87.67 | 87.67 | 12207 | 16.56 | 8039.4 |
| mapquik | 0.1 | 10 | 68418 | 26.31 | 38.46 | 42005 | 4.31 | 8466.6 |
| synpact | 0.1 | 10 | 99668 | 99.51 | 99.84 | 95 | 20.72 | 1449.5 |
| minimap2 | 0.1 | 15 | 100000 | 99.83 | 99.83 | 63 | 356.24 | 13323.3 |
| BLEND | 0.1 | 15 | 100000 | 85.36 | 85.36 | 14574 | 24.74 | 7790.6 |
| mapquik | 0.1 | 15 | 71845 | 26.30 | 36.61 | 45454 | 6.80 | 8516.6 |
| synpact | 0.1 | 15 | 99788 | 99.66 | 99.87 | 83 | 29.27 | 1456.5 |
| minimap2 | 0.1 | 20 | 100000 | 99.89 | 99.89 | 36 | 499.05 | 13320.2 |
| BLEND | 0.1 | 20 | 100000 | 83.28 | 83.28 | 16670 | 33.88 | 7984.1 |
| mapquik | 0.1 | 20 | 73506 | 26.42 | 35.94 | 47023 | 9.40 | 8146.6 |
| synpact | 0.1 | 20 | 99841 | 99.75 | 99.91 | 68 | 38.94 | 1479.4 |
| minimap2 | 0.1 | 25 | 100000 | 99.92 | 99.92 | 23 | 658.04 | 13316.1 |
| BLEND | 0.1 | 25 | 100000 | 81.62 | 81.62 | 18341 | 44.66 | 7716.9 |
| mapquik | 0.1 | 25 | 74307 | 26.60 | 35.79 | 47658 | 11.15 | 8178.4 |
| synpact | 0.1 | 25 | 99862 | 99.79 | 99.93 | 56 | 49.24 | 1491.0 |
| minimap2 | 0.5 | 10 | 100000 | 99.59 | 99.59 | 205 | 195.85 | 13308.9 |
| BLEND | 0.5 | 10 | 100000 | 88.76 | 88.76 | 10995 | 13.14 | 8183.8 |
| mapquik | 0.5 | 10 | 66516 | 26.00 | 39.09 | 40214 | 4.49 | 8142.6 |

continued on next page

*Table S5 (continued)*

| Mapper | Err. (%) | Len. (kb) | Mapped | Acc. (%) | Prec. (%) | W-chr | Time (s) | RSS (MB) |
|---|---|---|---|---|---|---|---|---|
| synpact | 0.5 | 10 | 99469 | 99.09 | 99.62 | 213 | 20.81 | 1445.6 |
| minimap2 | 0.5 | 15 | 100000 | 99.77 | 99.77 | 108 | 329.59 | 13314.4 |
| BLEND | 0.5 | 15 | 100000 | 86.58 | 86.58 | 13268 | 19.05 | 7895.0 |
| mapquik | 0.5 | 15 | 70409 | 26.10 | 37.07 | 44013 | 6.71 | 8136.6 |
| synpact | 0.5 | 15 | 99674 | 99.40 | 99.72 | 143 | 30.10 | 1448.8 |
| minimap2 | 0.5 | 20 | 100000 | 99.86 | 99.86 | 57 | 465.01 | 13311.5 |
| BLEND | 0.5 | 20 | 100000 | 84.63 | 84.63 | 15270 | 24.99 | 8124.4 |
| mapquik | 0.5 | 20 | 72142 | 26.42 | 36.62 | 45457 | 9.07 | 8148.6 |
| synpact | 0.5 | 20 | 99769 | 99.55 | 99.78 | 125 | 39.92 | 1454.7 |
| minimap2 | 0.5 | 25 | 100000 | 99.88 | 99.88 | 57 | 584.68 | 13315.1 |
| BLEND | 0.5 | 25 | 100000 | 83.00 | 83.00 | 16893 | 33.45 | 7836.7 |
| mapquik | 0.5 | 25 | 72686 | 26.49 | 36.44 | 46016 | 11.02 | 8294.6 |
| synpact | 0.5 | 25 | 99825 | 99.65 | 99.83 | 117 | 49.98 | 1461.7 |
| minimap2 | 1.0 | 10 | 100000 | 99.49 | 99.49 | 260 | 175.50 | 13308.9 |
| BLEND | 1.0 | 10 | 100000 | 90.01 | 90.01 | 9609 | 11.38 | 7954.4 |
| mapquik | 1.0 | 10 | 63703 | 25.46 | 39.97 | 37683 | 4.41 | 8672.6 |
| synpact | 1.0 | 10 | 98749 | 97.99 | 99.24 | 389 | 21.16 | 1435.9 |
| minimap2 | 1.0 | 15 | 100000 | 99.70 | 99.70 | 170 | 283.19 | 13311.0 |
| BLEND | 1.0 | 15 | 100000 | 88.17 | 88.17 | 11586 | 15.08 | 8123.4 |
| mapquik | 1.0 | 15 | 68142 | 25.89 | 37.99 | 41795 | 6.61 | 8298.5 |
| synpact | 1.0 | 15 | 99289 | 98.83 | 99.54 | 244 | 29.68 | 1436.9 |
| minimap2 | 1.0 | 20 | 100000 | 99.81 | 99.81 | 104 | 389.46 | 13307.9 |
| BLEND | 1.0 | 20 | 100000 | 86.48 | 86.48 | 13316 | 19.96 | 8138.8 |
| mapquik | 1.0 | 20 | 70081 | 26.00 | 37.10 | 43643 | 9.62 | 8726.6 |
| synpact | 1.0 | 20 | 99565 | 99.22 | 99.65 | 192 | 39.44 | 1440.6 |
| minimap2 | 1.0 | 25 | 100000 | 99.88 | 99.88 | 74 | 512.63 | 13307.9 |
| BLEND | 1.0 | 25 | 100000 | 84.89 | 84.89 | 14924 | 25.17 | 7717.9 |
| mapquik | 1.0 | 25 | 70351 | 26.08 | 37.07 | 43875 | 11.44 | 8162.7 |
| synpact | 1.0 | 25 | 99692 | 99.41 | 99.71 | 178 | 48.94 | 1446.9 |

Table S6: Complete real-data HiFi benchmark, evaluated by consensus of mappers. Acc. and Prec. are computed over reads in the consensus set; time excludes index loading.

| Genome | Mapper | Mapped | Acc. (%) | Prec. (%) | Map rate (%) | Time (s) | RSS (MB) |
|---|---|---|---|---|---|---|---|
| *Arabidopsis* | minimap2 | 99964 | 99.06 | 99.07 | 99.99 | 47.28 | 1375.2 |
| *Arabidopsis* | BLEND | 99903 | 97.30 | 97.34 | 99.93 | 15.30 | 1199.1 |
| *Arabidopsis* | mapquik | 95219 | 93.50 | 97.96 | 95.24 | 1.74 | 3651.0 |
| *Arabidopsis* | synpact | 99256 | 95.91 | 96.42 | 99.28 | 16.04 | 297.4 |
| Human | minimap2 | 99974 | 99.75 | 99.75 | 99.97 | 244.04 | 11857.9 |
| Human | BLEND | 99962 | 99.30 | 99.30 | 99.96 | 18.79 | 6814.7 |
| Human | mapquik | 99696 | 98.94 | 99.10 | 99.70 | 3.76 | 6346.6 |
| Human | synpact | 99793 | 99.04 | 99.11 | 99.79 | 28.99 | 815.0 |
| Maize | minimap2 | 99993 | 99.55 | 99.55 | 99.99 | 499.13 | 5767.2 |
| Maize | BLEND | 99931 | 98.76 | 98.79 | 99.93 | 19.97 | 4324.4 |
| Maize | mapquik | 98689 | 97.60 | 97.98 | 98.69 | 2.09 | 6895.5 |
| Maize | synpact | 99450 | 97.70 | 97.93 | 99.45 | 23.63 | 673.2 |
| Rye | minimap2 | 99998 | 99.48 | 99.48 | 100.00 | 430.65 | 13025.3 |
| Rye | BLEND | 99998 | 83.96 | 83.96 | 100.00 | 21.54 | 8011.8 |
| Rye | mapquik | 72694 | 26.11 | 35.83 | 72.69 | 10.20 | 9048.5 |
| Rye | synpact | 99565 | 97.34 | 97.54 | 99.56 | 31.50 | 1440.8 |

Table S7: Rye DeepConsensus real-data benchmark, evaluated by consensus of mappers. Acc. and Prec. are computed over reads in the consensus set; time excludes index loading.

| Genome | Mapper | Mapped | Acc. (%) | Prec. (%) | Map rate (%) | Time (s) | RSS (MB) |
|---|---|---|---|---|---|---|---|
| Rye | `minimap2` | 99999 | 99.92 | 99.92 | 100.00 | 366.03 | 13010.9 |
| Rye | BLEND | 99998 | 85.17 | 85.17 | 100.00 | 16.24 | 8001.5 |
| Rye | `mapquik` | 72527 | 27.20 | 37.52 | 72.53 | 5.79 | 9048.6 |
| Rye | `synpact` | 99663 | 99.58 | 99.75 | 99.66 | 30.92 | 1434.6 |
